# *XRN1* is a Species-Specific Virus Restriction Factor in Yeasts

**DOI:** 10.1101/069799

**Authors:** Paul A. Rowley, Brandon Ho, Sarah Bushong, Arlen Johnson, Sara L. Sawyer

## Abstract

In eukaryotes, the degradation of cellular mRNAs is accomplished by Xrn1p and the cytoplasmic exosome. Because viral RNAs often lack canonical caps or poly-A tails, they can also be vulnerable to degradation by these host exonucleases. Yeast lack sophisticated mechanisms of innate and adaptive immunity, but do use RNA degradation as an antiviral defense mechanism. One model is that the RNA of yeast viruses is subject to degradation simply as a side effect of the intrinsic exonuclease activity of proteins involved in RNA metabolism. Contrary to this model, we find a highly refined, species-specific relationship between Xrn1p and the “L-A” totiviruses of different *Saccharomyces* yeast species. We show that the gene *XRN1* has evolved rapidly under positive natural selection in *Saccharomyces* yeast, resulting in high levels of Xrn1p protein sequence divergence from one yeast species to the next. We also show that these sequence differences translate to differential interactions with the L-A virus, where Xrn1p from *S. cerevisiae* is most efficient at controlling the L-A virus that chronically infects *S. cerevisiae*, and Xrn1p from *S. kudriavzevii* is most efficient at controlling the L-A-like virus that we have discovered within *S. kudriavzevii*. All Xrn1p orthologs are equivalent in their interaction with another virus-like parasite, the Ty1 retrotransposon. Thus, the activity of Xrn1p against totiviruses is not simply an incidental consequence of the enzymatic activity of Xrn1p, but rather Xrn1p co-evolves with totiviruses to maintain its potent antiviral activity and limit viral propagation in *Saccharomyces* yeasts. Consistent with this, we demonstrated that Xrn1p physically interacts with the Gag protein encoded by the L-A virus, suggesting a host-virus interaction that is more complicated than just Xrn1p-mediated nucleolytic digestion of viral RNAs.

## Introduction

Degradation of mRNAs is a process essential to cell viability. Degradation pathways eliminate aberrant mRNAs, and also act to control gene expression levels. This process typically begins with host enzymes that perform either deadenylation or decapping on mRNAs targeted for degradation [1]. Following decapping, mRNAs are typically degraded by the 5’ to 3’ cytoplasmic exonuclease, Xrn1 [2,3]. Alternatively, after deadenylation, mRNAs can be subject to 3’ to 5’ degradation by the cytoplasmic exosome [4-6]. Viral transcripts and viral RNA genomes usually do not bear the canonical 5’ methylated cap structures or the 3’ polyadenylated (poly(A)) tails typical of cellular mRNAs, making them vulnerable to destruction by these host mRNA degradation pathways. In fact, it has been observed that Xrn1 and components of the exosome efficiently restrict virus replication in eukaryotes as diverse as mammals and yeasts [7-11]. As a result, mammalian viruses have evolved diverse countermeasures to prevent degradation by these proteins [7,8,12-18]. Still unknown is whether the host proteins like Xrn1 and components of the exosome can co-evolve with viruses to circumvent viral countermeasures. While such tit-for-tat evolution is common in mammalian innate immunity pathways, mRNA degradation is essential to the host and would be expected to be subject to strong evolutionary constraint.

*Saccharomyces* yeasts are known to harbor very few viruses [19]. Further, all yeast viruses are unable to escape their host cell, and instead are transmitted vertically through mating or during mitotic cell division. Almost all described species of *Saccharomyces* yeasts play host to double-stranded RNA (dsRNA) viruses of the family *Totiviridae*[20,21]. In fact, most commonly used *S. cerevisiae* laboratory strains are infected with a totivirus named L-A (Fig 1A) [22]. When initially synthesized, the RNAs produced by the L-A virus RNA-dependent RNA polymerase lack both a cap structure [23,24] and a poly(A) tail [25], and are vulnerable to degradation by Xrn1p [8] and the cytoplasmic exosome [14,26]. 3’-to-5’ degradation of viral RNAs by the cytoplasmic exosome is linked to the action of the SKI complex (Ski2, Ski3, Ski7, and Ski8), which acts to funnel aberrant RNAs into the nucleolytic core of the exosome [5,6]. The disruption of exosome and SKI complex genes has been shown to cause higher expression of viral RNAs, higher virus genome copy number, and an overproduction of virus-encoded toxins (i.e. the “superkiller” phenotype) [11,14,27]. In addition, the 5’-to-3’ exonuclease Xrn1p degrades viral transcripts and genomes of several RNA viruses in yeasts [8,24,28].

**Fig 1.**
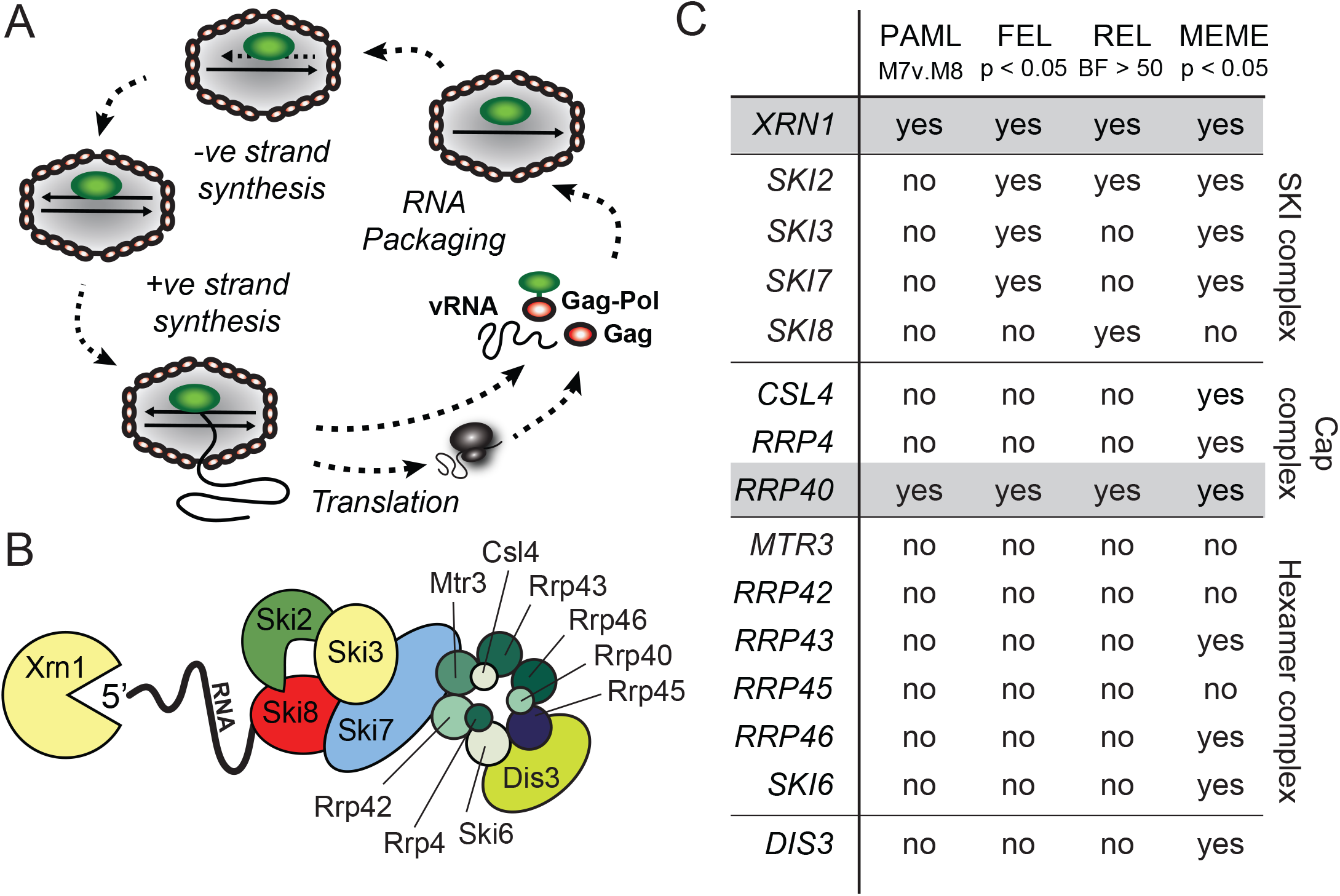
Evolutionary analysis of *Saccharomyces* genes involved in RNA metabolism. (A) Schematic of the lifecycle of the L-A virus of *S. cerevisiae*. Starting at the bottom of the figure, new viral positive sense single-stranded RNA (+ssRNA) is synthesized within the L-A virus capsid and extruded into the cytoplasm. The enzymatic activity of the viral Gag protein “steals” cap structures from host mRNAs and conjugates them to viral +ssRNAs. The capped viral +ssRNA is used as a template for translation and any remaining uncapped +ssRNA is encapsidated to form new viral particles by interaction with the L-A polymerase protein. Packaged +ssRNA is used as a template during negative strand synthesis to produce viral genomic dsRNA. (B) A cartoon representation of 5’-to-3’ and 3’-to-5’ RNA degradation by Xrn1p and the SKI/exosome complex, respectively, adapted from Parker *et al*. [29]. (C) Evolutionary analysis of *XRN1* and genes of the SKI/exosome complex. An alignment of each gene was analyzed using 4 common tests for positive selection (PAML, FEL, REL, and MEME) as fully described in Table S1. “Yes” indicates that there is positive selection detected in this gene by the indicated test.

Viruses and their hosts exist in a constant state of genetic conflict, where what is advantageous for one party is often disadvantageous for the other. Both genomes experience selection for mutations that benefit their own fitness but, particularly in yeast where viruses are strictly intracellular, the virus will be bounded in this process by the fact that if it begins to replicate too well, it may kill its host. Co-evolutionary battles between hosts and viruses play out in the physical interaction interfaces between interacting host and viral proteins (reviewed in [30-32]). One party is selected to reduce these interactions, and the other party is selected to strengthen them. For instance, there are many examples showing that mammalian restriction factors are selected to better recognize their viral targets, while viruses are continuously selected to escape that interaction, or to encode an antagonist protein that neutralizes the restriction factor. Because there is no equilibrium in these systems, this process cycles over and over, causing unusual signatures of evolution in both the host and virus proteins engaged in this interaction. While host protein complexes (host proteins interacting with other host proteins) can sometimes become co-evolved, this process of within-species refinement of protein-protein interactions is not the same as the dynamic and recurrent selection for new amino acids at interaction interfaces between host and pathogen proteins. The two scenarios can be disentangled using a metric that looks for codons that have accumulated a significantly higher rate of nonsynonymous mutations (dN) than even synonymous mutations (dS). The signature of dN/dS > 1 commonly results from the repeated cycles of selection that occurs in genetic conflict scenarios [43], but has never been shown to be driven by subtler processes like the refinement of within-host physical interactions.

Host genes in genetic conflict with viruses diverge in sequence in a manner that alters interactions with viruses. For this reason, these proteins become non-equivalent between host species from the perspective of viruses. Highly diverged host proteins reinforce species barriers, making it difficult for viruses to move from their current host species into new host species (for example, [33,34]). Since yeast have limited antiviral strategies, we reasoned that evolutionary pressure on the RNA quality control pathways to thwart the replication of RNA viruses might be especially intense. This led us to investigate the unique evolutionary scenario involving a restriction system employing proteins critical to RNA turnover and cellular homeostasis.

In this study, we analyzed whether or not any components of the yeast RNA degradation pathways mentioned above are evolving under positive natural selection, potentially indicative of tit-for-tat coevolution with viruses. We identified this evolutionary signature in at least two genes involved in RNA metabolism, *RRP40* and *XRN1*, and then undertook an in depth functional analysis of *XRN1*. To test the hypothesis that Xrn1p has been honed by co-evolution to target and restrict totiviruses, we made a series of *S. cerevisiae* strains where *XRN1* is replaced with wild-type orthologs from other *Saccharomyces* species. All *XRN1* orthologs fully complemented an *XRN1* knockout strain of *S. cerevisiae*, as assessed by several assays. On the other hand, we found that *XRN1* orthologs were different in their ability to control the replication of the L-A virus. Xrn1p from *S. cerevisiae* was most efficient at controlling the L-A virus that chronically infects *S. cerevisiae*, and Xrn1p from *S. kudriavzevii* was most efficient at controlling the L-A-like virus (SkV-L-A1) that we discovered within *S. kudriavzevii*. All *XRN1* orthologs were equivalent in their interaction with another virus-like parasite, the Ty1 retrotransposon. Our identification of signatures of positive selection and species-specific virus restriction suggests that *XRN1* can be tuned by natural selection to better restrict totivirus in response to the evolution of these viruses over time. We show that the structure of Xrn1p affords the flexibility to change in response to selective pressure from totiviruses, while also maintaining cellular functions.

## Results

### Components of the yeast RNA degradation pathway are evolving under positive selection

To differentiate between models where Xrn1p restricts L-A through a passive mechanism that is incidental to its inherant exonuclease activity, or through an active mechanism where Xrn1p evolves to optimally suppress L-A replication, we first looked for evidence of positive selection (dN/dS > 1) within the genes encoding the major components of the SKI complex, the exosome, and Xrn1p (Fig 1B). Importantly, signatures of positive selection do not identify the genes that are most important for controlling viral replication. These statistical tests are designed to identify host proteins that are involved in direct physical interactions with viruses, and which also have the evolutionary flexibility to change in response to viral selective pressure, becoming species-specific in the process. For this reason, we would neither expect to identify signatures of positive selection in all genes known to be involved in controlling totiviruses, nor in all genes encoding components of a complex like the exosome. For each gene, we collected sequences from six divergent species of *Saccharomyces* (*S. cerevisiae*, *S. paradoxus*, *S. mikatae*, *S. kudriavzevii*, *S. arboricolus*, and *S. bayanus*) [44-46] and created a multiple sequence alignment. We then analyzed each alignment for evidence of codons with dN/dS > 1 using four commonly employed tests for positive selection [47,48]. We see some evidence for positive selection of specific codon sites in several of these genes, however, only *XRN1* and the exosome subunit gene *RRP40* passed all four tests (Fig 1C; Table S1). Other genes are determined to be under positive selection by some tests, and may be of interest to explore further. Of *XRN1* and *RRP40*, the impact of *XRN1* on viral replication has been more directly substantiated [7-9,13-16,35-37,49-53], so we focused our attention on this gene. However, it should be noted that *RRP40* encodes a component of the cytoplasmic exosome, which, in conjunction with the SKI complex, is clearly linked to the restriction of L-A [11,14,27].

### Xrn1p is species-specific in the restriction of L-A virus

We next tested if *S. cerevisiae XRN1* has been tailored by co-evolution with the L-A virus. Double-stranded RNA (dsRNA) purified from an *S. cerevisiae xrn1* strain migrates as a distinct band of 4.6 kilobase pairs (Fig 2A), which is consistent with the size of the L-A virus genome, and its identity was further confirmed by RT-PCR (Fig S1). We confirmed a strong reduction in dsRNA when the *xrn1* strain was complemented with plasmid-mounted *XRN1* from *S. cerevisiae* under the transcriptional control of its native promoter (Fig 2A), consistent with the published role of Xrn1p as an L-A restriction factor [8,14,24,27]. On the other hand, catalytically-dead versions of Xrn1p (E176G and Δ1206-1528) did not suppress L-A dsRNA levels (Fig 2B), as has been previously described [54]. We next performed heterospecific (other species) complementation by introducing the *XRN1* from *S. mikatae*, *S. kudriavzevii*,or *S. bayanus* into the *S. cerevisiae xrn1Δ* strain. These species were chosen as they are representative of the diversity found within the *sensu stricto* complex of *Saccharomyces* yeasts. Strikingly, no other Xrn1p was able to reduce L-A dsRNA to the same extent as Xrn1p from *S. cerevisiae* (Fig 2A). Xrn1p from *S. mikatae*, the closest relative to *S. cerevisiae* in this species set, was capable of slightly reducing L-A dsRNA abundance. Xrn1p from *S. bayanus* and *S. kudriavzevii* appear to have levels of dsRNA similar to *xrn1Δ*, indicating little or no effect on L-A copy number. In summary, we find that *XRN1* orthologs vary in their ability to restrict the *S. cerevisiae* L-A virus. This is somewhat surprising for a critical and conserved gene involved in RNA quality control, but consistent with the signatures of positive selection which suggest that certain parts of this protein are highly divergent between species.

**Fig 2.**
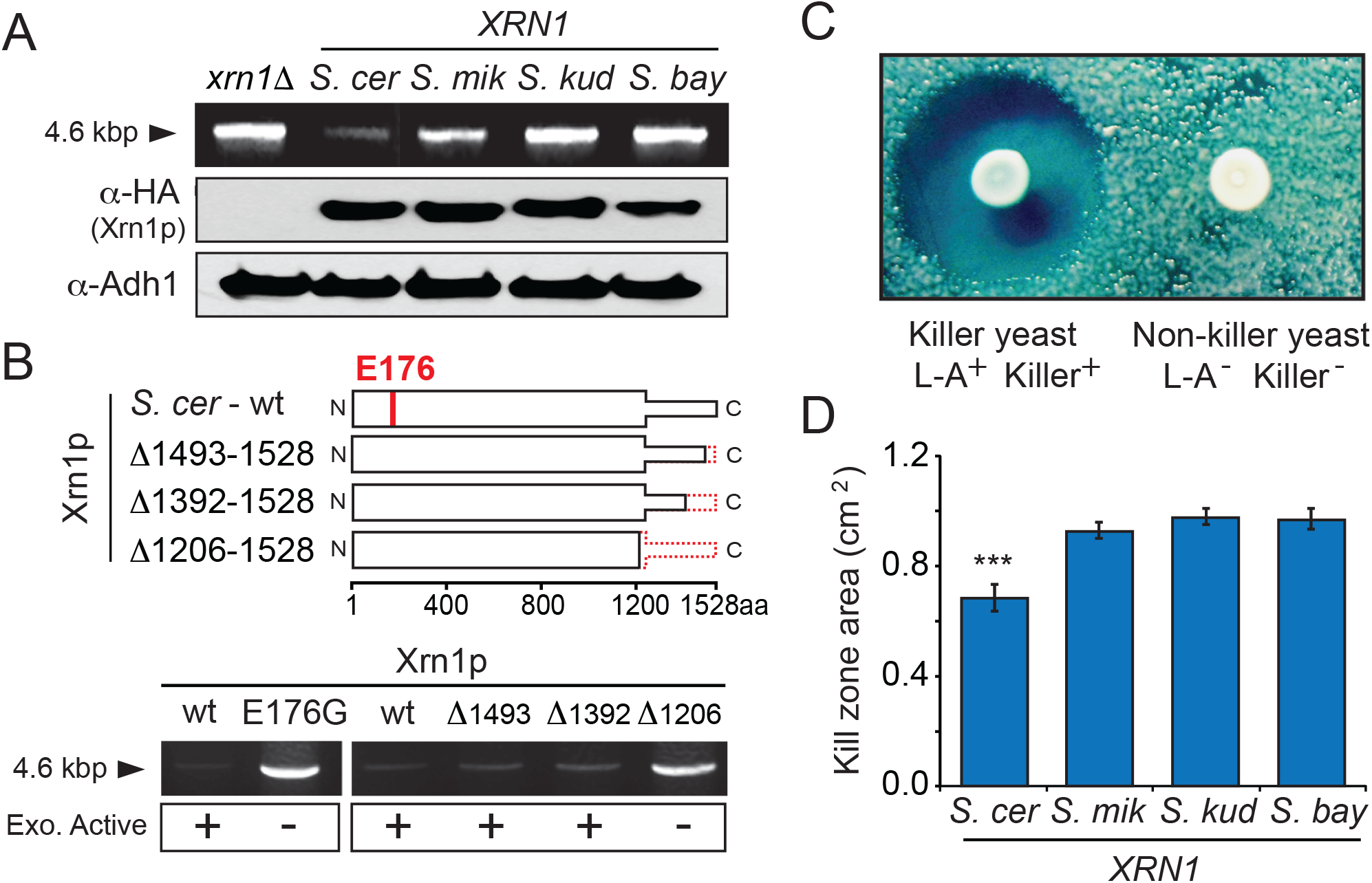
The effect of *XRN1* evolution on the restriction of the L-A and killer viruses of *S. cerevisiae*. (A) (*Top row*) dsRNA extraction from *S. cerevisiae xrn1Δ* with or without complementation by *XRN1* from different species of *Saccharomyces* (*S. cer – S. cerevisiae*; *S. mik – S. mikatae*; *S. kud – S. kudriavzevii; S. bay – S. bayanus*). From the same agarose gel, the first two lanes were spliced to the last three lanes for clarity. (*Bottom two rows*) Western blots showing the expression of HA-tagged Xrn1 and Adh1 (control) within each complemented strain. (B) (*Top*) Domain diagrams of Xrn1p showing the position of the catalytic residue E176 and the location of the C-terminal truncations of Xrn1p (dashed line). Xrn1p with the E176G mutation or the Δ1206-1528 truncation are catalytically inactive, as described previously [54]. (*Bottom*) dsRNA extraction from *S. cerevisiae xrn1* expressing wild type *XRN1*, the catalytically inactive *XRN1*(E176G) or truncation mutants of Xrn1p (all derived from *S. cerevisiae*). (C) A representative picture of *S. cerevisiae* killer (L-A^+^ Killer^+^) and non-killer (L-A^-^ Killer ^-^) yeasts and the effect of killer toxin expression on the growth of a lawn of sensitive yeast. (D) The effect of Xrn1p expression on the diameter of the kill zones around killer yeast. Asterisks are indicative of a significant difference in mean kill zone area compared to all other samples tested (Tukey-Kramer test p<0.05). See figure S2 for example images of the kill zones for each sample.

We next used a functional and quantitative assay to confirm the species-specific effects of *XRN1* on virus replication. This assay exploits the dsRNA “killer virus” (also known as M virus). The killer virus is a satellite RNA of L-A and is totally dependent on L-A proteins for replication. It uses L-A-encoded proteins to encapsidate and replicate its genome, and to synthesize and cap its RNA transcripts [12]. The killer virus encodes only a single protein, a secreted toxin referred to as the killer toxin [19,55,56]. The result is that “killer yeast” colonies, i.e. those infected with both L-A and the killer virus, kill neighboring cells via the diffusion of toxin into the surrounding medium (Fig 2C). Importantly, resistance to the killer toxin is provided by the pre-processed, immature form of the toxin, supplying killer yeast cells with an antidote to their own poison [55]. It has been shown previously that Xrn1p can inhibit the expression of the killer phenotype by degrading uncapped killer virus RNAs [14,57]. Therefore, we use the presence and size of kill zones produced by killer yeasts as a quantitative measurement of killer virus RNA production in the presence of each Xrn1p ortholog.

A strain of *S. cerevisiae* lacking *XRN1*, but harboring both the L-A and killer virus (*xrn1* L-A^+^ Killer^+^), was complemented with each *XRN1* ortholog. Clonal isolates from each complemented strain were grown to mid-log phase, and 6 x 10^5^ cells were spotted onto an agar plate seeded with a lawn of toxin-sensitive yeast. After several days’ incubation at room temperature, kill zones around these culture spots were measured and the total area calculated. The transformation of *xrn1* L-A^+^ Killer^+^ with *S. cerevisiae XRN1* produced an average kill zone that covered 0.68 cm^2^ (n = 14). However, transformation with *XRN1* from *S. mikatae*, *S. bayanus*, or *S. kudriavzevii* produced significantly larger kill zones covering 0.92 cm^2^ (n = 11), 0.96 cm^2^(n = 17) and 0.97 cm^2^ (n = 17), respectively. The kill zone produced by *xrn1Δ* L-A^+^ Killer^+^ yeast expressing *S. cerevisiae XRN1* was significantly smaller than those produced by yeast expressing any of the other *XRN1* ortholog (Tukey–Kramer test, p<0.05) (Figs 2D). The smaller kill zones in the strain expressing *S. cerevisiae XRN1* are consistent with lower levels of killer and L-A derived RNAs. In summary, this assay also supports a species-specific restriction phenotype for *XRN1*.

It has been observed that over-expression of *XRN1* can cure *S. cerevisiae* of the L-A virus, presumably by degrading viral RNA so effectively that the virus is driven to extinction [8,28]. Therefore, we developed a third assay to test the ability of *XRN1* orthologs to control L-A, in this case by assessing their ability to cure *S. cerevisiae* of the virus. Plasmids expressing HA-tagged and untagged Xrn1p were transformed into a killer strain of *S. cerevisiae* with its genomic copy of *XRN1* intact. This was followed by the analysis of more than 100 purified clones for virus curing, that is, the absence of the killer phenotype as indicated by the loss of a kill zone when plated on a lawn of sensitive yeast. Importantly, the introduction of an empty plasmid fails to produce any cured clones (n = 103) (Figs 3A and 3B). Provision of an additional copy of *S. cerevisiae XRN1* cured 49% of clones (n = 159) (Figs 3A and 3B). Cured clones remained cured (i.e. non-killers) when purified and tested again for their ability to kill sensitive yeasts (n = 20). Over-expression of *XRN1* from *S. mikatae*, *S. kudriavzevii*, and *S. bayanus* was unable to efficiently cure the killer phenotype, resulting in only 12% (n = 129), 8% (n = 120), and 9% (n = 123) cured clones, respectively (Fig 3A, blue bars). The loss of L-A from cured strains was also verified by RT-PCR. We detected no L-A or killer RNAs within the four cured clones analyzed (Fig 3C). These data show that *XRN1* from all *Saccharomyces* species have the ability to cure the killer phenotype, however, *XRN1* from *S. mikatae*, *S. kudriavzevii*, and *S. bayanus* is considerably less efficient than *S. cerevisiae XRN1*. Taken together, we show that viral restriction by *XRN1* is species-specific. These data contradict a model where viral restriction is merely incidental to the RNA quality control functions of *XRN1*, but is rather something that can be refined through sequence evolution in *XRN1*.

**Fig 3.**
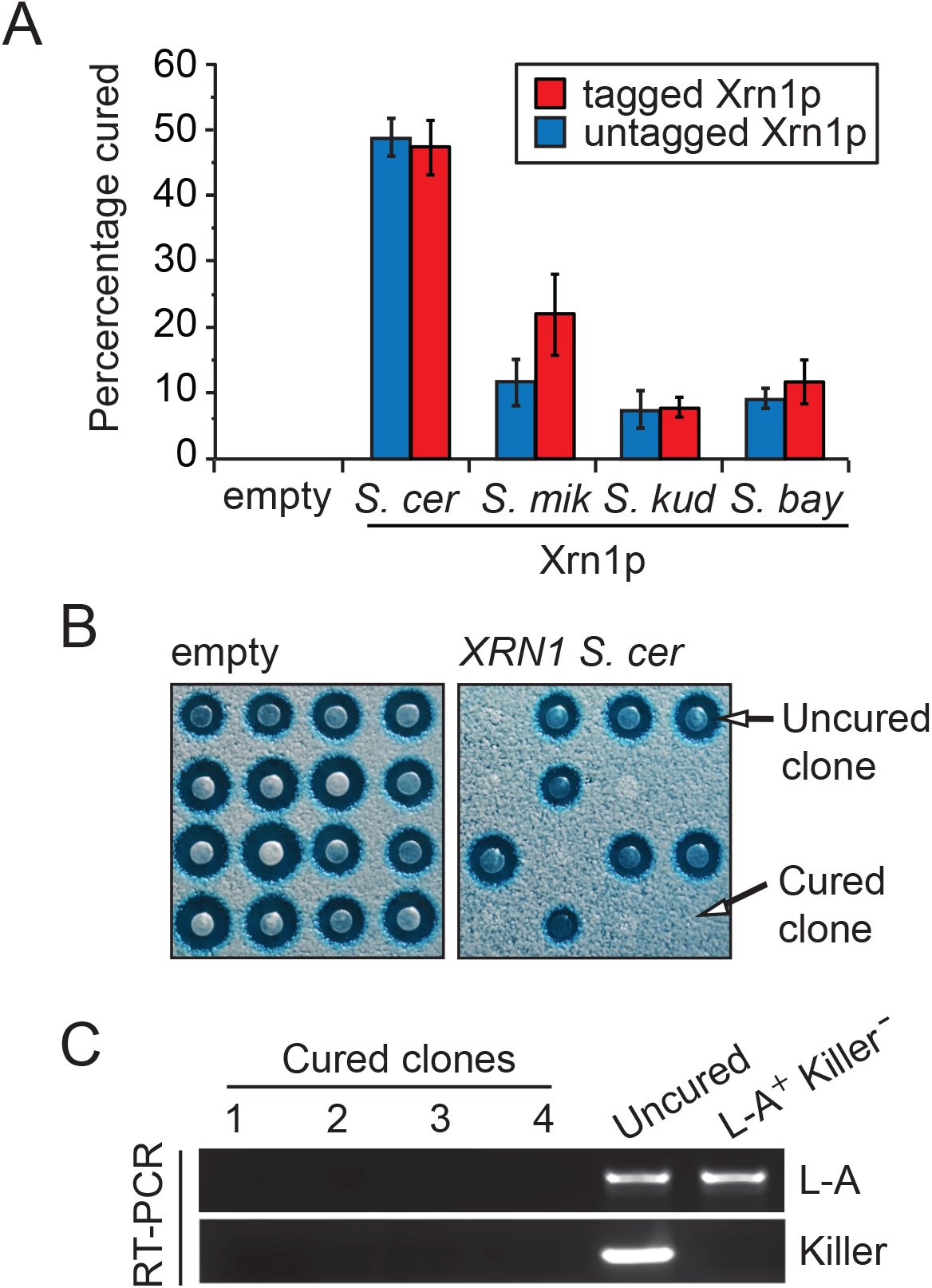
*XRN1* orthologs vary in their ability to cure *S. cerevisiae* of the L-A and killer viruses. (A) Killer *S. cerevisiae* strains over-expressing Xrn1p orthologs (with or without C-terminal HA tag) were assayed for loss of the killer phenotype resulting in “cured” clones. The percentage of cured clones is indicated. (B) Representative data from 16 clonal isolates either not expressing (*left*) or expressing *XRN1* from *S. cerevisiae* (*right*), with the lack of kill zone indicating a cured clone. (C) Non-quantitative RT-PCR was used to confirm the loss of L-A and killer RNAs from strains deemed to have lost the killer phenotype from Fig 3A and 3B, compared to the parental uncured strain (L-A^+^ Killer^+^) and *S. cerevisiae* BY4741 (L-A^+^ Killer^-^).

### *XRN1* cellular function has been conserved in *Saccharomyces* species

We next tested the presumption that the *XRN1* orthologs are functionally equivalent for cellular processes when expressed within *S. cerevisiae*. We first confirmed that *XRN1* orthologs successfully complemented the severe growth defect of *S. cerevisiae xrn1Δ*, by measuring the doubling time of *S. cerevisiae xrn1* with or without a complementing *XRN1Δ*-containing plasmid (Fig 4A). The knockout of *XRN1* also renders cells sensitive to the microtubule-destabilizing fungicide benomyl [54], and we observed that all *XRN1* homologs convey equal resistance to benomyl on solid medium (Fig 4B). It has been previously reported that over-expression of *XRN1* is toxic to *S. cerevisiae*, a phenotype that has been suggested to be due to a dominant negative interaction of Xrn1p with other essential cellular components, such as the decapping complex [54]. Growth upon medium containing 2% galactose was equivalently reduced for strains carrying *GAL1* inducible *XRN1* genes from each species, whereas the strain over-expressing *GFP* grew normally (Fig 4C, right). Finally, the Ty1 retrotransposon is another intracellular virus that replicates within *Saccharomyces* species and often co-exists with L-A within the same cell. Interestingly, Xrn1p is not a restriction factor for Ty1, but rather promotes Ty1 replication [35-42]. We found no significant difference between the mean values for retrotransposition in the presence of Xrn1p from *S. cerevisiae*, *S. mikatae, S. kudriavzevii*, or *S. bayanus* (one-way ANOVA, F_3, 8_=0.36, p=0.78), indicating that the evolutionary differences between divergent *XRN1* genes do not affect the ability of Ty1 to replicate within *S. cerevisiae* (Fig 4D). Collectively, these data indicate the cellular functions of Xrn1p have remained unaffected during yeast speciation, while the interaction with L-A viruses has changed.

**Fig 4.**
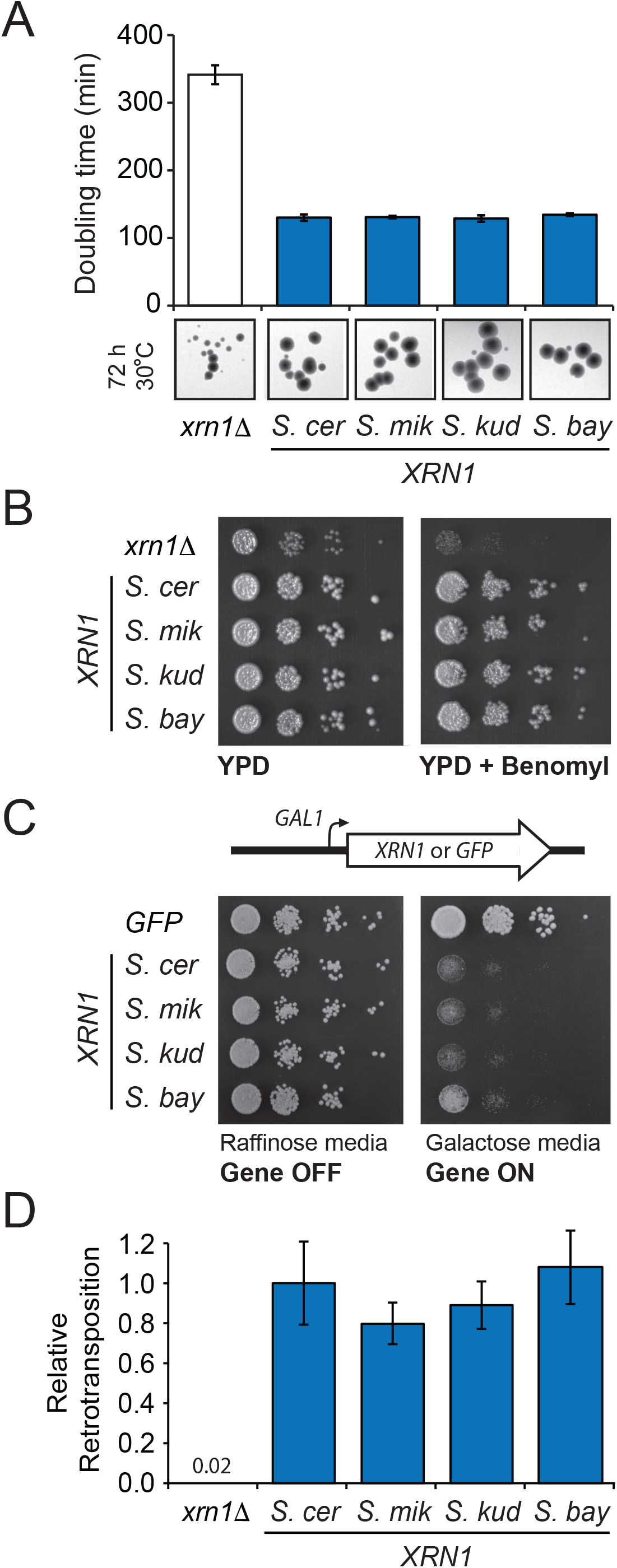
*XRN1* has conserved its housekeeping functions during *Saccharomyces* speciation. (A) The doubling time of *S. cerevisiae xrn1* with and without complementation of *XRN1Δ* from different species of *Saccharomyces* (error bars represent SEM, n = 3). Below are representative pictures of the growth and morphology of individual colonies. (B) The growth and benomyl sensitivity of *S. cerevisiae xrn1* cultured on YPD solid medium, and the effect of complementation with *XRN1Δ* from different species of *Saccharomyces*. (C) The effect of over-expression of each *XRN1Δ* ortholog on the growth of *S. cerevisiae* upon solid medium, compared to the over-expression of *GFP.* Cells are grown on medium containing either raffinose or galactose as the sole carbon source to control the activity of the *GAL1* promoter. (D) Ty1 retrotransposition within *S. cerevisiae xrn1Δ* complemented by *XRN1* from different species, relative to *XRN1* from *S. cerevisiae*. All error bars represent SEM, n>3.

### Signatures of positive selection in *XRN1* correlate with a region important for L-A antagonism

We next mapped the region responsible for the species-specific restriction by *XRN1*. To better understand the structural organization of Xrn1p from *S. cerevisiae*, we used Phyre [58] to generate a template-based homology model of the exonuclease using the solved structure of *Kluyveromyces lactis* Xrn1p (Fig 5A). A linker region within the N-terminal domain, the far C-terminal domain, and domain D2 were not included in the model as there is a lack of information regarding the structural organization of these regions. Importantly, modeled portions contained three of the residue positions that we identified as evolving under positive selection (Table S1) and all of these (blue) fall in and near the D1 domain (orange) (Fig 5A). As expected because of the selection that has operated on them, these residue positions under positive selection are more variable in sequence between species than are surrounding residues (two are shown in Fig 5B). All residues under positive selection are surface exposed and are far from the highly conserved Xrn1p catalytic domain (96% identity, across the *Saccharomyces* genus) and catalytic pocket (red). The other sites of positive selection fall within the last 500 amino acids of Xrn1p, which is less conserved compared to the rest of the protein (83% identity, across the *Saccharomyces* genus) (Fig. 5C).

**Fig 5.**
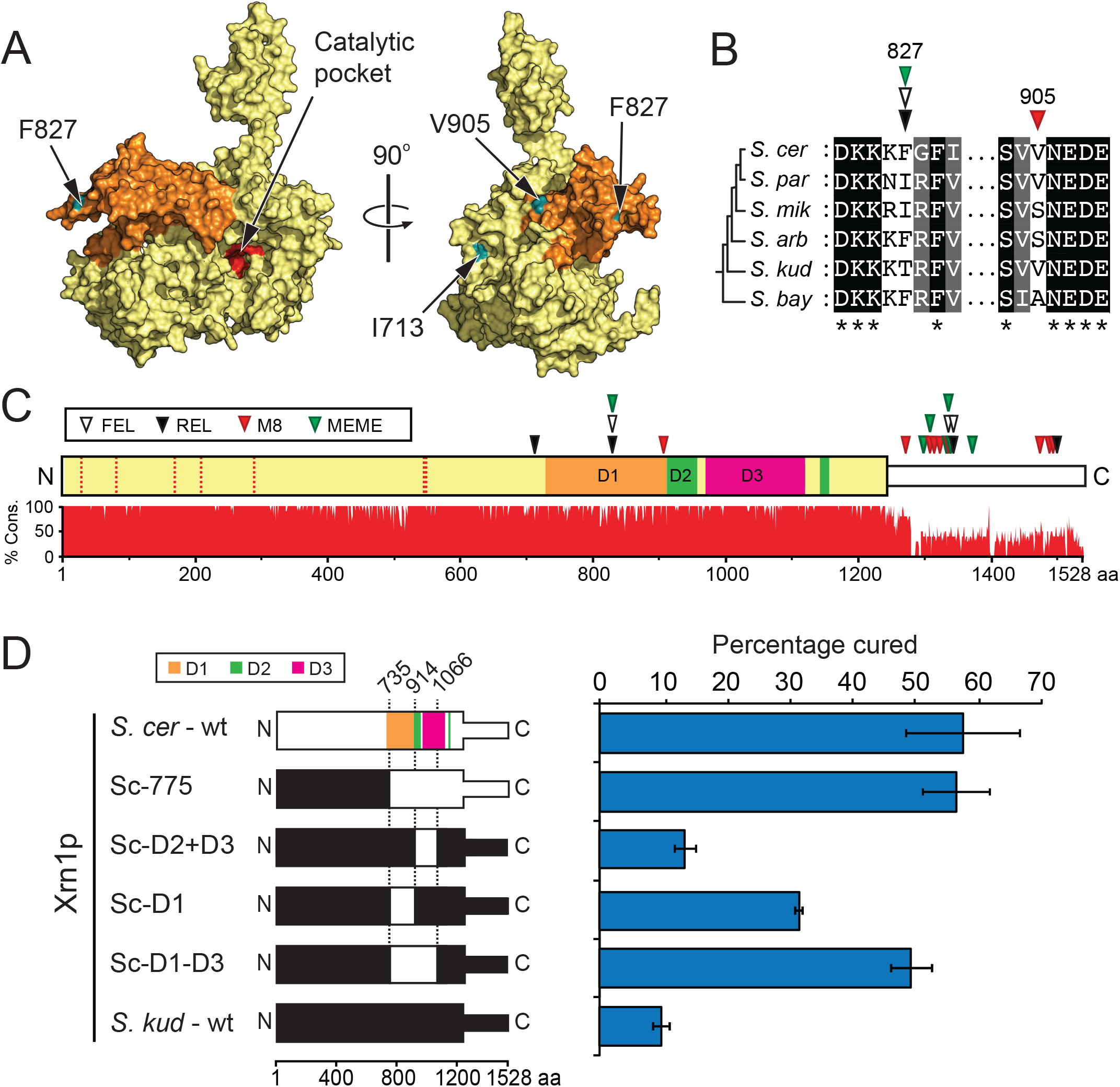
A structured protein domain within Xrn1p is responsible for species-specific antiviral activity. (A) Space-filled structural model of *S. cerevisiae* Xrn1p generated by Phyre analysis [58]. The D1 domain is colored orange. Amino acids 354–503, 979–1109, and 1240-1528 are unresolved in the model due to a lack of structural information. The structural model is included as File S1. (B) A representative amino acid alignment shows two variable sites within the Xrn1p D1 domain that are predicted to be evolving under positive selection. (C) A linear diagram of *S. cerevisiae* Xrn1p based on the structure of *K. lactis* Xrn1p, indicating select domains of the C-terminus and showing the conservation of Xrn1p across *Saccharomyces* species. The white domain has an unknown structure and is predicted to be unstructured by Phyre analysis [58]. Triangles indicate the position of sites deemed to be under positive selection by four site-based models of molecular evolution, PAML M8 (red), REL (black), FEL (white), and MEME (green). See Table S1 for a list of all sites. The seven residues that form the catalytic pocket of Xrn1p are shown as vertical dotted lines within the N-terminal domain. (D) (*Left*) Chimeric proteins derived from the fusion of domains of Xrn1p from *S. cerevisiae* (white) and *S. kudriavzevii* (black). Relevant domains within Xrn1p are colored according to Chang *et al.* (D1 (orange) 731-914; D2 (green), 915-960 and 1134-1151; D3 (magenta); 978-1108) [59]. (*Right*) Clonal isolates of a killer *S. cerevisiae* strain expressing chimeric Xrn1p were assayed for their ability to cure the killer phenotype from *S. cerevisiae* (error bars represent SEM, n>3).

To define the importance of the two regions that we identified as containing signatures of positive selection, we replaced portions of *S. kudriavzevii XRN1* with the equivalent portions of *S. cerevisiae XRN1*, and assayed for a region of *S. cerevisiae XRN1* that would convey the ability to cure the killer phenotype. We found that an *XRN1* chimera encoding the last 775 amino acids from *S. cerevisiae* (Sc-775) was sufficient to cure 56% of clones analyzed, and this was very similar to *S. cerevisiae XRN1* (57%) (Fig 5D). Conversely, when the last 777 amino acids from *S. kudriavzevii* (Sk-777) were used to replace the same region within *S. cerevisiae XRN1*, only 9% of clones were cured (Fig S3). This focused our construction of further chimeras to the second half of the protein, which also contains all of the codons under positive selection and has less amino acid conservation between *S. cerevisiae* and *S. kudriavzevii* (82% protein identity, compared to the N-termini of Xrn1p with 95% identity).

Initial analysis of the highly diverged C-terminal tail revealed that the last 461 amino acids of *S. cerevisiae* Xrn1p were unable to convey efficient L-A restriction to *S. kudriavzevii* Xrn1p (Fig S3). For this reason, we focused further chimeric analysis on the region encompassing the D1-D3 domains (Fig 5A-C), as defined previously [59]. We swapped into *S. kudriavzevii* Xrn1p the D2-D3, D1, or D1-D3 domains of *S. cerevisiae* Xrn1p, and saw increasing rescue of the ability to cure the L-A virus (Fig 5D). All chimeric *XRN1* genes were functionally equivalent with respect to their cellular functions, as all were able to establish normal growth and benomyl resistance in *S. cerevisiae xrn1* (Fig S4). Therefore, the species-specificity domain maps predominantly to D1, with contribution from the neighboring D2 and D3 domains as well. Together, our data suggest that the exonuclease activity of Xrn1p is important for virus restriction and is preserved across species, but that evolution has tailored a novel virus interaction domain (D1-D3) that targets the enzymatic activity of Xrn1p against L-A in a manner that changes over time, keeping step with virus evolution.

### Xrn1p physically interacts with the L-A virus Gag proteins

It’s hard to imagine that Xrn1p proteins from different species are differentially recognizing viral RNA, since they all retain their host functionalities in RNA degradation. We considered the possibility that there might be host-virus interactions beyond Xrn1p and the viral RNA. It has been shown that Xrn1p targets uncapped viral RNA transcripts rather than affecting dsRNA propagation [57]. As totivirus transcription only occurs in the context of a fully-formed capsid [60] and capsids are assembled entirely from the L-A Gag protein [61], it would seem plausible that Xrn1p may interact directly with Gag to target virus-derived uncapped RNAs. We introduced epitope tags onto Xrn1p (HA-tag) and the major capsid protein of L-A, Gag (V5-tag), and expressed both tagged and untagged versions of each protein from plasmids introduced into *S. cerevisiae xrn1*Δ (Fig 6A). Bead-bound antibodies specific for either HA or V5 were used to immunoprecipitate Xrn1p or Gag, respectively. We found that Gag (V5-tagged) was able to immunoprecipitate Xrn1p (HA-tagged) from *S. cerevisiae* and *S. kudriavzevii* (Fig 6A, top panel). Reciprocally, Xrn1p-HA from both *S. cerevisiae* and *S. kudriavzevii* were able to immunoprecipitate Gag-V5 (Fig 6A, bottom panel). The interaction between Xrn1p and Gag appears not to be mediated by single-stranded RNAs, as their digestion by RNase A in the whole cell extract did not affect the co-immunoprecipitation of Gag by Xrn1p (Fig S5). We next performed these experiments with a monoclonal antibody specific to L-A Gag, so that endogenous L-A Gag protein could be immunoprecipitated. This reaction co-immunoprecipitated both *S. cerevisiae* and *S. kudriavzevii* Xrn1p (Fig 6B). Qualitatively, the relative efficiencies of Gag interaction with both *S. cerevisiae* and *S. kudriavzevii* Xrn1p appear similar in all assays, which seems at odds with our model that suggests that evolutionary differences within Xrn1p are a direct determinant of totivirus interaction. There are several possible interpretations. First, Gag might be antagonizing Xrn1p rather than being the species-specific target of Xrn1. Second, there may be a third component in this interaction which makes manifest the species-specificity of Xrn1p. Finally, a trivial explanation could be that coimmunoprecipitations are not very quantitative, and maybe there is in fact a difference in interaction with Gag between the Xrn1p of different species. Nonetheless, these data demonstrate a previously undescribed interaction that goes beyond Xrn1p interaction with viral RNA and warrants careful *in vitro* study.

**Fig 6.**
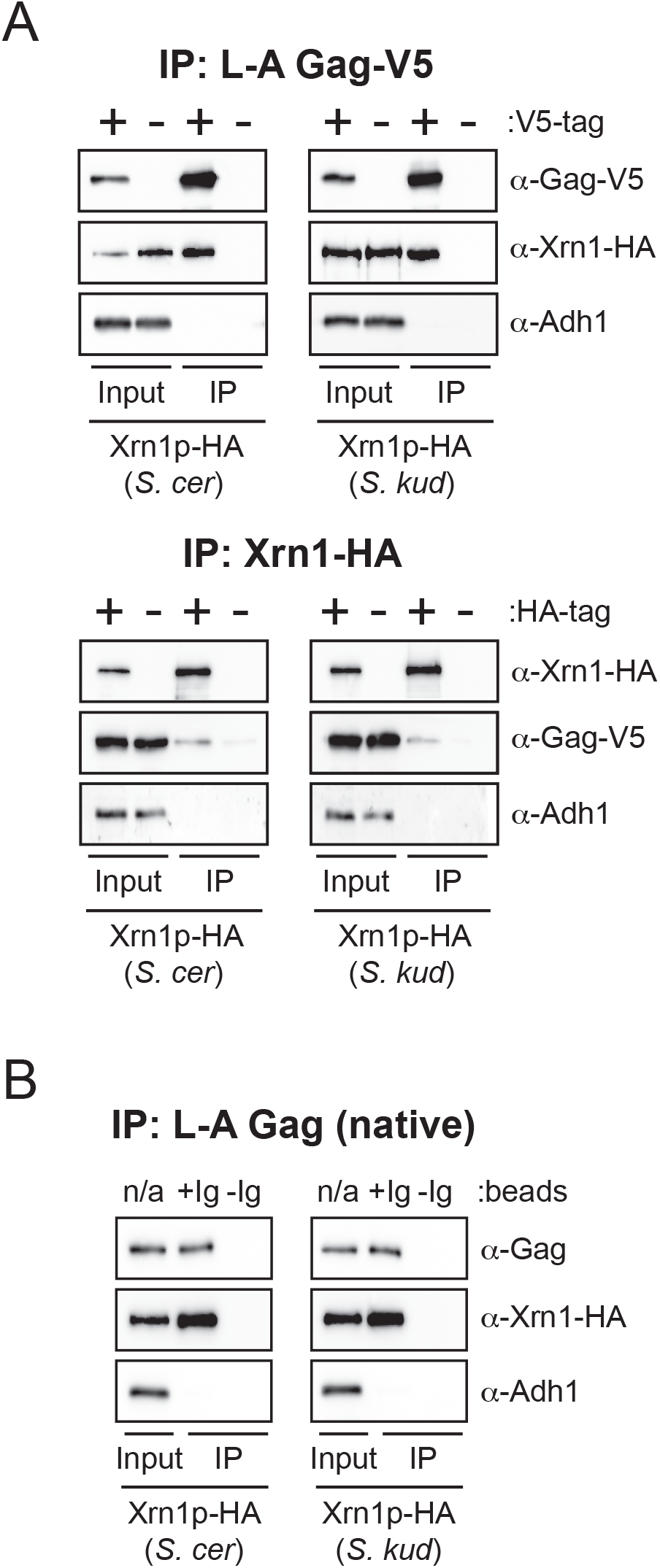
The totivirus structural protein Gag is associated with Xrn1p *in vivo*. Western blot analysis of Xrn1p and L-A Gag co-immunoprecipitation. (A) (*Top*) V5-tagged and untagged Gag proteins were immunoprecipitated in the presence of Xrn1p-HA from either *S. cerevisiae* or *S. kudriavzevii*. (*Bottom*) HA-tagged and untagged Xrn1p from either *S. cerevisiae* or *S. kudriavzevii* were immunoprecipitated in the presence of Gag-V5. (B) Native, untagged, virus encoded L-A Gag was immunoprecipitated in the presence of Xrn1p-HA from either *S. cerevisiae* or *S. kudriavzevii* using beads with (+Ig) or without (-Ig) anti-Gag antibody present. Adh1p was used in all panels as a loading control to ensure equal input of total protein and the specificity of immunoprecipitation.

### Confirmation of the species-specific restriction of Xrn1p using a newly described totivirus from *S. kudriavzevii.*

We next wished to test our findings against other related yeast viruses. Indeed, the *S. cerevisiae* totivirus L-A-lus has been shown to have limited susceptibility to *XRN1* from a different strain of *S. cerevisiae* [28]. We also wanted to test viruses of other species, but the only fully characterized totiviruses within the *Saccharomyces* genus are from *S. cerevisiae*. To identify totiviruses of other species, we screened *Saccharomyces* species from the *sensu stricto* complex for the presence of high molecular weight viral RNA species, and discovered a ~4.6 kbp dsRNA molecule within *S. kudriavzevii* FM1183 isolated from Europe (Fig 7A) [46]. We cloned the 4.6 kbp dsRNA molecule using techniques described by Potgieter *et al*. [62] and sequenced the genome of the virus using Sanger sequencing. We named the virus SkV-L-A1 (*S*.*kudriavzevii* virus L-Aisolate number 1; Genbank accession number: KX601068). The SkV-L-A1 genome was found to be 4580 bp in length, with two open reading frames encoding the structural protein Gag and the fusion protein Gag-Pol (via a-1 frameshift) (Fig 7B). Conserved features of totiviruses were identified and include the conserved catalytic histidine residue required for cap-snatching (H154), a-1 frameshift region, packaging signal, and replication signal (Figs 7B and S6). Phylogenetic analysis of the Gag and Pol nucleotide and protein sequences firmly places SkV-L-A1 within the clade of *Saccharomyces* totiviruses represented by L-A and L-A-lus [28,63], as opposed to the more distantly related *Saccharomyces* totivirus L-BC (Fig 7C) [64].

**Fig 7.**
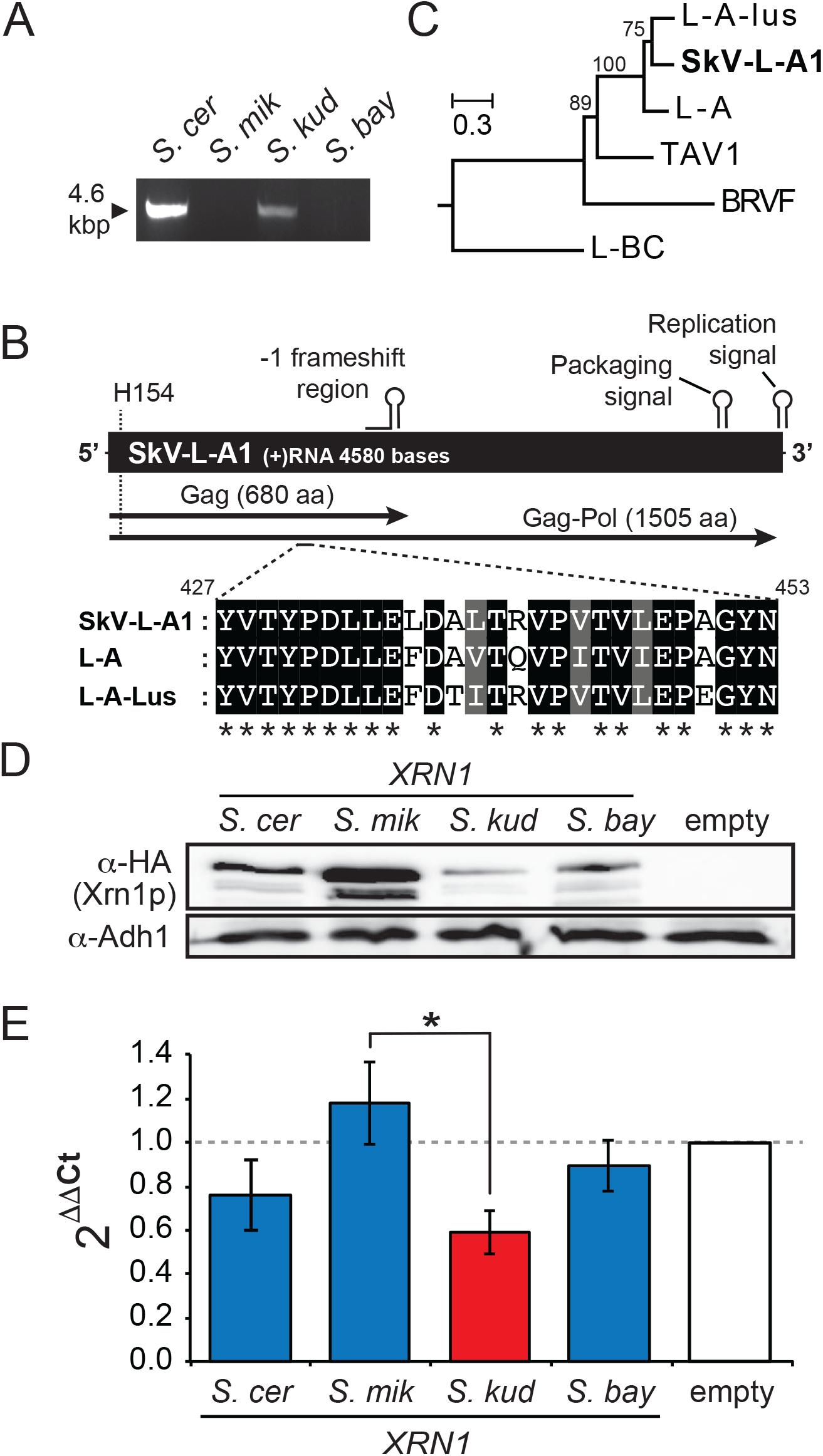
The description of a novel totivirus from *S. kudriavzevii* and its unique sensitivity to restriction by Xrn1p from its cognate host species. (A) dsRNA extraction from different species of *Saccharomyces* yeasts including *S. cerevisiae* (BY4741), *S. mikatae* (JRY9181), *S. kudriavzevii* (FM1183), and *S. bayanus* (JRY8153). The major product of the extraction method is a single ~4.6 kbp dsRNA species, as shown by agarose gel electrophoresis. (B) A schematic representation of the genome organization of the totivirus SkV-L-A1 from *S. kudriavzevii*. H154 represents the conserved catalytic histidine used for totivirus cap-snatching [14]. (C) The evolutionary relationship of SkV-L-A1 to known totiviruses was inferred by using the Maximum Likelihood method with bootstrap values from 100 replicates shown at each node. The nucleotide sequence of the *POL* gene was aligned from six totiviruses (GenBank accession numbers: SkV-L-A1 (this study; KX601068), L-A-lus (JN819511), L-A (NC_003745), tuber aestivum virus 1 (TAV1) (HQ158596), black raspberry virus F (BRVF) (NC_009890), L-BC (NC_001641)). The tree is drawn to scale, with branch lengths measured in the number of substitutions per site. The exact phylogenetic relationship of the “L-A-like” viruses is somewhat ambiguous due to low bootstrap support within the clade (Fig S6). (D) *S. kudriavzevii* was used to express HA-tagged *XRN1* from different *Saccharomyces* yeasts (*S. cer – S. cerevisiae*; *S. mik – S. mikatae*; *S. kud – S. kudriavzevii; S. bay – S. bayanus*), with their expression measured by Western blotting, relative to the expression of *ADH1*. (E) Relative abundance of SkV-L-A1 RNAs when *XRN1* from different *Saccharomyces* species are over-expressed within *S. kudriavzevii*, as determined by RT-qPCR. The asterisk represents a p-value of <0.05 (Tukey-Kramer test).

To test the effect of *XRN1* upon SkV-L-A1, plasmids expressing *XRN1* orthologs were introduced via LiAc transformation into *S. kudriavzevii* infected with SkV-L-A1. These plasmids were able to express *XRN1* from each species, although we find that the expression is variable with *S. mikatae* Xrn1p expressing at the a level higher than the others (Fig 7D). The expression of these proteins did not affect the overall growth rate or colony morphology of *S. kudriavzevii* (Fig. S7). Because of the lack of an observable killer phenotype in this strain (likely because a killer toxin-encoding satellite dsRNA is not present), heterospecific *XRN1* were expressed within *S. kudriavzevii* and analyzed for their ability to spontaneously cure SkV-L-A1, as we did previously with L-A in *S. cerevisiae* (Fig 3). We did not observe any virus curing by any orthologs of Xrn1p, but believe that this could be because the high-copy plasmids that we used in this experiment in *S. cerevisiae* are unable to drive Xrn1p expression in *S. kudriavzevii* high enough to actually cure the virus. However, we have observed previously that Xrn1p can reduce the abundance of totivirus RNAs (Fig 2A), so we further analyzed the *XRN1*-transformed clones of *S. kudriavzevii* for changes in SkV-L-A1 RNA levels using reverse transcriptase quantitative PCR (RT-qPCR). Total RNA was extracted from clones of *S. kudriavzevii* and converted to cDNA using random hexamer priming. cDNA samples were amplified using primers designed to specifically target SkV-L-A1 *GAG* and the cellular gene *TAF10*. The empty vector control was used as the calibrator sample, and *TAF10* expression was used as the normalizer to calculate the relative amount of SkV-L-A1 RNAs present within each *XRN1* expressing *S. kudriavzevii* cell line using the comparative *C*T method [65]. We found that expression of *XRN1* from *S. kudriavzevii* (n = 10) reduced the relative levels of SkV-L-A1 RNAs by 40% (Fig 7E), even though this Xrn1p was expressed at the lowest levels (Fig 7D). This is in contrast to *XRN1* from *S. mikatae* (n = 9) and *S. bayanus* (n = 8) that only showed a 13% increase or 15% decrease in SkV-L-A1 RNAs, respectively. *S. cerevisiae XRN1* was able to reduce SkV-\L-A1 RNAs by 27% and is noteworthy due to the close evolutionary relationship between SkV-L-A1 and other L-A-like viruses from *S. cerevisiae* (Fig S6). These data suggest that Xrn1p is a species-specific restriction factor in different *Saccharomyces* yeasts, and that coevolution of totiviruses and yeasts has specifically tailored the potency of Xrn1p to control the replication of resident viruses within the same species.

## Discussion

In the *Saccharomyces* genus, Xrn1p, the SKI complex, and exosome are all important for controlling the abundance of totivirus RNAs. We find that *XRN1* and the exosome component *RRP40* are somewhat unique in their strong signatures of positive natural selection. We speculated that positive selection might be driven by selection imposed by totiviruses. As speciation occurs and viruses mutate in unique ways in each lineage, new allelic versions of these antiviral genes that enable better control of totivirus replication would experience positive natural selection. Indeed, we found this to be the case, with *S. cerevisiae* Xrn1p restricting the *S. cerevisiae* L-A virus better than any other ortholog of *XRN1*, and *S. kudriavzevii* Xrn1p restricting *S. kudriavzevii* SkV-L-A1 virus the best. The exact nature of the host-virus protein-protein interaction that is driving this evolutionary arms race is not clear. To thwart *XRN1*, the totiviruses are known to synthesize uncapped RNAs with an exposed 5’ diphosphate, which is a suboptimal substrate for Xrn1p-mediated decay [24]. Further, it has been shown that the totivirus Gag protein has a cap-snatching activity that cleaves off caps from host mRNAs and uses them to cap viral transcripts, protecting them from Xrn1p degradation [12,14]. We have found that Xrn1p interacts with L-A Gag, and that this interaction is not mediated by the presence of single-stranded RNAs. What remains unknown is whether Xrn1p is targeting Gag as part of the restriction mechanism, or whether Gag is targeting Xrn1p as a counter defense. As we did not observe an obvious species-specific differences in the interaction between Xrn1p and L-A Gag by coimmunoprecipitation, we cannot clearly define the observed role of sequence variation in Xrn1p. This may be because of the low sensitivity of our assay system, or because direct binding of Xrn1p by L-A Gag is ubiquitous and that the rapid evolution of *XRN1* results from another intriguing facet of virus-host interaction and antagonism. However, we now know that the interaction between L-A and Xrn1p goes beyond the simple recognition of L-A RNA by Xrn1p. We can speculate that Xrn1p may compete with Gag for access to uncapped viral RNAs as they are extruded into the cytoplasm, or that interaction with unassembled Gag allows the recruitment of Xrn1p to sites of virion assembly resulting in viral RNA degradation. Alternately, it is possible that the target of Xrn1p is simply L-A RNA, and that the interaction with Gag reflects a viral countermeasure where Gag is redirecting or otherwise altering the availability of Xrn1p to degrade L-A RNA. Indeed, there are several examples of mammalian viruses that redirect or degrade Xrn1p to aid in their replication [17,18,66].

The literature suggests that Xrn1p is a widely-utilized restriction factor against viruses, as it has been reported to have activity against mammalian viruses [9,16], yeast viruses [8,24], and plant viruses [53]. The potent 5’-3’ exonuclease activity of Xrn1p has resulted in viruses developing a rich diversity of strategies to protect their RNAs. For instance, Hepatitis C virus recruits MiR-122 and Ago2 to its 5’ UTR to protect its RNA genome from Xrn1p degradation [7,16]. The yeast single-stranded RNA narnavirus uses a different strategy to protect its 5’ terminus, folding its RNA to form a stem-loop structure that prevents Xrn1p degradation [8]. In some cases, viruses even depend on Xrn1p to digest viral RNA in a way that benefits viral replication, for example, preventing the activation of innate immune sensors [49]. Flavivirus (West Nile and Dengue virus) genomes also encode RNA pseudoknot and stem-loop structures that arrest the processive exonuclease activity of Xrn1p, producing short subgenomic flavivirus RNAs (sfRNAs) that are important for viral pathogenicity [13,67]. Members of the *Flaviviridae*, *Herpesviridae*, *Coronaviridae*, and yeast *Totiviridae* have all been shown to encode proteins that initiate endonucleolytic cleavage of host mRNAs, revealing exposed 5’ monophosphates that are substrates for Xrn1p degradation. This is thought to interfere with host translation and to produce uncapped RNA “decoys” that potentially redirect Xrn1p-mediated degradation away from viral RNA [14,15]. Xrn1p degradation, Xrn1p relocalization, virus-encoded capping enzymes, cap-snatching mechanisms, RNA-protein conjugation, recruitment of host micro-RNAs, cleavage of host mRNAs as “decoys”, and viral RNA pseudoknots are all utilized to prevent Xrn1p-mediated viral RNA destruction [7,8,12-18]. All of this evidence suggests that viruses can employ various methods to escape or harness the destructive effects of Xrn1. Our data now suggests that Xrn1p in yeast is not a passive player in the battle against viruses, but rather that hosts can be selected to encode new forms of Xrn1p that can overcome viral strategies.

To rationalize the model of an antagonistic relationship between L-A and *Saccharomyces* species, it is important to consider the fitness burden of strictly intracellular viruses. Prevailing wisdom assumes that infection of fungi by viruses is largely asymptomatic and benign, especially when considering that their intracellular lifecycle ensures an evolutionary dead-end if they kill or make their host unfit. Indeed, within laboratory yeast strains, the association between L-A and *S. cerevisiae* appears to be at equilibrium, with no major biological differences between strains infected or not infected by L-A [68]. Therefore, the relationship between L-A and the *Saccharomyces* yeasts could be viewed as mutualistic or even commensalistic [68-70]. Mutualism is particularly striking in the context of the L-A / killer virus duo that provides the host cell with the “killer” phenotype, a characteristic that is broadly distributed throughout fungi [71]. If an infected yeast cell can kill other yeasts around it using the killer toxin, it no longer has to compete for resources within that environmental niche, an evolutionarily advantageous situation [56,69,70]. Indeed, there are other examples of host-virus mutualism in fungi [72,73]. However, there are many observations that lead one to believe that the relationship between intracellular viruses and their hosts is not benign and static. Firstly, there is a measurable fitness cost to killer toxin production by *S. cerevisiae* within unfavorable environmental conditions that inactivate the toxin, allow for regular cellular dispersal and/or are nutrient rich [69,70]. Secondly, virus infection of pathogenic fungi can also cause hypovirulence (a reduction in fungal pathogenesis), an outcome that is being exploited to treat agricultural disease [74-77]. Thirdly, many wild and domesticated strains of *S. cerevisiae* are free of totiviruses (and therefore also of killer), suggesting that there is selection against the ongoing maintenance of these viruses [20,28,71]. Fourthly, the continued maintenance of RNAi systems in fungi also correlates with the loss of the killer phenotype and is known to antagonize fungal viruses [71,78]. However, a virus of the fungi *Cryphonectria parasitica* has been shown to antagonize and escape restriction by RNAi without crippling its host [78]. This antagonistic relationship appears similar to the equilibrium of *Saccharomyces* yeasts and totiviruses, and suggests that in the absence of effective RNAi, additional antiviral defenses may be biologically relevant (i.e. Xrn1p). In line with this view of a dynamic relationship between hosts and intracellular viruses, we show that totiviruses from different *Saccharomyces* species are best controlled by the Xrn1p of their cognate species, and that disruption of this equilibrium can result in excessive virus replication (Fig 2), virus loss (Fig 3), or a reduction in viral RNA (Fig 2 and Fig 7). Signatures of positive selection that we have detected in *Saccharomyces XRN1* are also consistent with a host-virus equilibrium that is in constant flux due to the dynamics of a back-and-forth evolutionary conflict (Fig 2 and Fig 6).

There are several examples of mammalian housekeeping proteins engaged in evolutionary arms races with viruses. (By “housekeeping” we refer to proteins making critical contributions to host cellular processes, as opposed to proteins dedicated to immunity.) In most of these other examples though, the housekeeping protein is hijacked by viruses to assist their replication in the cell (rather than serving to block viral replication). For instance, many viruses hijack cell surface receptors to enter cells. We and others have shown that entry receptors are quite evolutionarily plastic, and that mutations can reduce virus entry without compromising host-beneficial functions of the receptor [33,79-83]. For example, the antagonistic interaction of Ebola virus (and/or related filoviruses) with the bat cell surface receptor, Niemann-Pick disease, type C1 (NPC1), has driven the rapid evolution of the receptor without affecting the transport of cholesterol, critical to the health of the host [33]. Numerous such examples highlight how essential housekeeping machineries, not just the immune system, are critical for protecting the cell from replicating viruses. This study highlights an interesting evolutionary conundrum that does not apply to classical immunity genes: as Xrn1p appears to be an antiviral protein, it must be able to evolve new antiviral specificities without compromising cellular health and homeostasis.

## Materials and Methods

### Plasmid construction

*XRN1* from *S. cerevisiae*, including 1000 bp of the 5’ and 3’ UTRs, was amplified by PCR from genomic DNA prepared from *S. cerevisiae* S288C. This PCR product was cloned into the plasmid pAG425-GAL-*ccd*B by the “yeast plasmid construction by homologous recombination” method (recombineering) [84] to produce pPAR219. Briefly, pAG425-GAL-*ccd*B was amplified by PCR to produce a 5000 bp product lacking the *GAL-1* gene and the *ccd*B cassette. The PCR primers used to amplify pAG425-GAL-*ccd*B contained additional DNA sequence with homology to the UTRs of *XRN1* from *S. cerevisiae*. Both PCR products were used to transform BY4741, with correctly assembled plasmids selected for by growth on complete medium (CM) –leucine. The *XRN1* open reading frame (HA-tagged and untagged) from *S. mikatae, S. bayanus*,or *S. kudriavzevii* was introduced into pPAR219 between the 5’ and 3’ UTRs from *S. cerevisiae XRN1* using recombineering to produce pPAR225, pPAR226, and pPAR227, respectively. As a negative control, *NUP133* was cloned into the pPAR219 plasmid backbone to produce pPAR221, which was used to allow growth of *xrn1* on medium lacking leucine without *XRN1* complementation. The *LEU2* gene was replaced by *TRP1* using recombineering techniques to produce the plasmids pPAR326, pPAR327, pPAR328, and pPAR329. Using PCR and recombineering, we also constructed chimeric *XRN1* genes by exchanging regions of *S. kudriavzevii XRN1* (pPAR227) with the corresponding regions of *S. cerevisiae XRN1* (pPAR219). *XRN1* inducible plasmids were constructed by cloning PCR-derived *XRN1* genes into pCR8 by TOPO-TA cloning (Thermo Fisher). Utilizing Gateway^TM^ technology (Thermo Fisher), *XRN1* genes were sub-cloned into the destination vector pAG426-GAL-*ccd*B for over-expression studies [85]. The same pCR8/Gateway workflow was also used to clone and tag *GAG* from a cDNA copy of the L-A totivirus (pI2L2) to produce pPAR330 and pPAR331. The DNA sequences from all constructed plasmids can be found in File S2.

### Yeast strain construction

The *S. cerevisiae* killer strain (BJH001) was created by the formation of a heterokaryon from the mating of the haploid strains BY4733 (*KAR1*) and 1368 (*kar1*) [86]. The resultant daughter heteroplasmon cells were selected by growth on CM –uracil and the ability to produce zones of growth inhibition indicative of the presence of L-A and the killer virus. The inability to grow on CM lacking histidine, leucine, tryptophan or methionine was also used to confirm the genotype of BJH001. BJH006 was created by replacing *XRN1* with the *KANMX4* gene using homologous recombination within BJH001 [87].

### Rapid extraction of viral dsRNA from *Saccharomyces cerevisiae*

1 × 10^9^ yeast cells (~10 mL) were harvested from a 24-48 h overnight culture grown to saturation. Strains of *S. kudriavzevii* and *S. mikatae* were grown at room temperature, all other strains were grown at 30°C. The flocculent nature of some strains of wild yeasts made it challenging to accurately determine the exact number of cells present in some cultures. In these cases, the size of the cell pellet was used as an approximate measure of cell number relative to *S. cerevisiae*. Harvested cells were washed with ddH_2_O, pelleted, and washed with 1 ml of 50 mM EDTA (pH 7.5). Cells were again harvested and the pellets suspended by vortexing in 1 ml of 50 mM TRIS-H_2_SO_4_ (pH9.3), 1% β-mercaptoethanol (added fresh), and incubated at room temperature for 15 min. The cell suspension was centrifuged and the supernatant removed and the cell pellet suspended in 1 ml of BiooPure-MP (a single-phase RNA extraction reagent containing guanidinium thiocyanate and phenol) (Bioo Scientific) and vortexed vigorously. 200 μl of chloroform was added and vortexed vigorously before incubation for 5 min at room temperature. The aqueous phase and solvent phase were separated by centrifugation at 16,000 x g for 15 min at 4°C. The aqueous phase was transferred to a new tube and 1/3 volume of 95-100% ethanol added and mixed well by vortexing. The entire sample was loaded onto a silica filter spin column (Qiagen plasmid miniprep kit) and centrifuged for 30 s at 16,000 × g. The flow-through was discarded and the column washed twice with 750 μl of 100 mM Nacl/75% ethanol by centrifugation at 16,000 × g for 30 sec. The column was dried by centrifugation at 16,000 × g for an additional 30 sec. The dsRNA was eluted from the column by the addition of 100 μl of 0.15 mM EDTA (pH 7.0) and incubation at 65°C for 5 min before centrifugation at 16,000 × g for 30 sec.

### Detection of L-A by RT-PCR

dsRNA that was extracted from 1 × 10^9^ yeast cells using our rapid extraction of viral dsRNA protocol was used as template for superscript two-step RT-PCR (Thermo Fisher). cDNA was created using a primer specific for the negative strand L-A genomic RNA – 5’ CTCGTCAGCGTCTTGAACAGTAAGC. Primers 5’-GACGTCCCGTACCTAGATGTTAGGC and 5’-CTCGTCAGCGTCTTGAACAGTAAGC were used to specifically target and amplify cDNA derived from negative strand L-A virus RNAs using PCR with Taq (New England Biolabs). The plasmid pI2L2 was used as a positive control for the RT-PCR reaction as it contains a cDNA copy of the L-A virus genome [88]. Alternatively, we collected total RNA from ~1 × 107 actively growing yeast cells using the RNeasy total RNA extraction kit (Qiagen) and synthesized cDNA using primers to target both the positive and negative strand of either L-A (5’-AAGATATTCGGAGTTGGTGATGACG and 5’-TCTCCGAAATTTTTCCAGACTTTATAAGC) or killer virus (5’-GCGATGCAGGTGTAGTAATCTTTGG and 5’-AGTAGAAATGTCACGACGAGCAACG). The same primers were used to detect L-A and killer virus specific cDNAs using PCR with Taq polymerase (New England Biolabs).

### Ty1 retrotransposition assays

We assayed Ty1 retrotransposition in *S. cerevisiae xrn1Δ*, using the previously described Ty1 retrotransposition reporter system [89], and confirmed that *XRN1* deletion causes a dramatic reduction in Ty1 retrotransposition (~50-fold) [35]. To test the effect of *XRN1* evolution on Ty1 replication, we introduced *XRN1* from *S. cerevisiae*, *S. mikatae, S. kudriavzevii*, or *S. bayanus* into *xrn1*Δ and assayed Ty1 retrotransposition.

### Western blot Analysis of Xrn1p

Yeast lysates were prepared using the Y-PER reagent (Thermo Fisher) from 100 μl volume of log-phase yeast cells as per manufacturers instructions or by bead beating as described previously [90]. HA-tagged Xrn1p was detected via Western blot using a 1:5000 dilution of a horseradish peroxidase conjugated anti-HA monoclonal antibody (3F10 - # 12013819001) (Roche). Adh1p was detected using a 1:10000 dilution of rabbit polyclonal anti-alcohol dehydrogenase antibody Ab34680 (Abcam). V5-tagged proteins were detected using a 1:5000 dilution of a mouse monoclonal antibody (R960-25) (Life Technologies). Native L-A Gag was detected using a 1:1000 dilution of a mouse monoclonal antibody (gift from Nahum Sonenberg). Secondary antibodies were detected using ECL Prime Western Blotting Detection Reagent on a GE system ImageQuant LAS 4000 (GE Healthcare Life Sciences).

### Evolutionary analysis

Nucleotide sequences from six species of *Saccharomyces* yeasts were obtained from various online resources, where available [44,46,91]. Maximum likelihood analysis of dN/dS was performed with codeml in the PAML 4.1 software package [47]. Multiple protein sequence alignments were created using tools available from the EMBL (EMBOSS Transeq and Clustal Omega) (www.embl.de). Protein alignments were manually curated to remove ambiguities before processing with PAL2NAL to produce accurate DNA alignments [92]. DNA alignments were fit to the NSsites models M7 (neutral model of evolution, codon values of dN/dS fit to a beta distribution, with dN/dS > 1 not allowed) and M8 (positive selection model of evolution, a similar model to M7 but with an additional site class of dN/dS > 1 included in the model). To ensure robustness of the analysis, two models of codon frequencies (F61 and F3×4) and multiple seed values for dN/dS (ω) were used (Table S1). Likelihood ratio tests were performed to evaluate which model of evolution the data fit significantly better. Posterior probabilities of codons under positive selection within the site class of dN/dS > 1 (M8 model of positive selection) were then deduced using the Bayes Empirical Bayes (BEB) algorithm. REL and FEL analysis was carried out using the online version of the Hyphy package (www.datamonkey.org) Table S1 [48]. Analysis of *XRN1* was performed using the TrN93 nucleotide substitution model and the following phylogenetic relationship (Newick format):

((((((*S. paradoxus*-Europe, *S. paradoxus*-Far East), (*S. paradoxus*-North America, *S. paradoxus*-Hawaii)), *S. cerevisiae*), *S. mikatae*), *S. kudriavzevii*), *S. arboricolus*, *S. bayanus*);

GARD analysis found no significant evidence of homologous recombination within any dataset. MEGA6 was used to infer the evolutionary history of totiviruses using the Maximum Likelihood method. Appropriate substitution models were selected using manually curated DNA and protein alignments. The tree topologies with the highest log likelihood were calculated, with all positions within the alignment files containing gaps and missing data ignored. The reliability of the generated tree topologies was assessed using the bootstrap test of phylogeny using 100 iterations. Bootstrap values >50% are shown above their corresponding branches.

### Benomyl sensitivity assay

YPD plates containing 15 μg ml^-1^ of benomyl were prepared as described previously [54]. Yeast strains expressing *XRN1* or containing an empty vector were grown overnight at 30°C in CM –leucine. Cell numbers were normalized and subject to a 10-fold serial dilution before spotting onto YPD agar plates with or without benomyl, and grown at 37°C for 72 h.

### Over-expression of *XRN1* and the effect on cell growth

*S. cerevisiae* carrying multi-copy plasmids encoding *XRN1* or *GFP* under the control of the *GAL1* promoter were grown overnight at 30°C in CM –uracil with raffinose as a carbon source. Cell numbers were normalized and subject to a 10-fold serial dilution before spotting onto CM –uracil agar plates containing either 2% raffinose or galactose. Plates were grown at 30°C for 72 h.

### Kill zone measurement

Plasmids encoding various *XRN1* genes were used to transform BJH006. Purified single colonies of killer yeasts were inoculated in 2 ml CM –leucine cultures and grown to mid-log phase. YPD “killer assay” agar plate supplemented with methylene blue (final concentration 0.003% w/v) and pH balanced to 4.2 with sodium citrate, were freshly inoculated and spread with *S. cerevisiae* K12 and allowed to dry. Thereafter, 1.5 μl of water containing 6 × 10^5^ cells was spotted onto the seeded YPD plates and incubated at room temperature for 72 h. The diameter of the zones of growth inhibition were measured and used to calculate the total area of growth inhibition.

### Killer phenotype curing assay

The curing of the killer phenotype was measured by transforming *S. cerevisiae* BJH006 with approximately 100 ng of plasmid encoding various *XRN1* genes using the LiAc method. The addition of 1000 ng or as little as 10 ng of plasmid had no affect on the percentage of colonies cured using this assay. After 48 h of growth, colonies were streaked out and grown for a further 48 h. Clonal isolates of killer yeasts were patched onto a YPD “killer assay” plate (see kill zone measurement protocol) that were previously inoculated with *S. cerevisiae* K12, and incubated at room temperature for 72 h. The presence or absence of a zone of inhibition was used to calculate the percentage of killer yeast clones cured of the killer phenotype.

### Xrn1p structural modeling

PHYRE was used to create a template-based homology model of *S. cerevisiae* Xrn1p using the solved structure of *K. lactis* Xrn1p as a template [58,59]. The structure was determined with an overall confidence of 100% (36% of aligned residues have a perfect alignment confidence as determined by the PHYRE inspector tool), a total coverage of 81%, and an amino acid identity of 67% compared to *K. lactis* Xrn1p. PDB coordinates for the modeled structure can be found in File S1. Structural diagrams were constructed using MacPyMOL v7.2.3.

### Co-immunoprecipitation of Xrn1p and L-A Gag

Strains were grown in CM lacking the appropriate amino acids in order to retain the relevant plasmids. For co-immunoprecipitations involving L-A Gag-V5 and Xrn1p-HA, 50 mL cultures (CM –tryptophan –leucine, 2% raffinose) were used to inoculate 500 mL cultures (CM –tryptophan –leucine, 2% galactose) at OD_600_~0.1. Cells were harvested at OD_600_0.7, after ~14 h of growth at 30°C with shaking. Cultures used for the immunoprecipitation of native Gag were grown in the same manner, but in CM –leucine medium containing 2% dextrose. Immunoprecipitation of yeast and viral proteins were performed as previously described [90] with the following modifications: 2-4 mg of protein was used per co-immunoprecipitation. Approximately 50 μg of protein was loaded for the whole-cell extract “input”, as determined by Bradford Assay (~2% of total input), and was compared to 10-20% of each co-immunoprecipitation. Sepharose beads were substituted for Dynabeads® MyOne^TM^ Streptavidin T1 or Dynabeads® Protein G (Thermo Fisher Scientific). For immunoprecipitation of Xrn1p-HA, we used an anti-HA-Biotin, rat monoclonal antibody (3F10 -#12158167001) (Roche), and for Gag-V5 a mouse monoclonal antibody (R960-25) (Life Technologies). RNase A was added to whole cell extracts at a concentration of (80 µg mL^-1^) and incubated with Dynabeads during immunoprecipitation for 2 hours at 4°C. RNAse is in excess in our co-IP experiments, because significant RNA degradation occurred at concentrations of RNase 8-fold lower than we used (Figure S5). RNA from samples with and without the addition of RNase A was recovered from yeast whole cell extracts after co-immunoprecipitation using Trizol according to manufacturer’s guidelines (Thermo Fisher). The extent of RNA degradation was measured using a 2200 TapeStation Instrument and a RNA screentape, as per manufacturer’s instructions (Agilent). An RNA integrity number (RIN) was calculated for each sample based upon criteria that reflect the quality of the RNA sample, as described previously [93].

### Cloning and sequencing of SkV-L-A1

dsRNAs were isolated from *S. kudriavzevii* as described above and processed according to the protocol of Potgieter *et al.* [62], with the following modifications: Reverse transcription reactions were carried out using Superscript IV (Thermo Fisher), PCR amplification was performed by Phusion polymerase (Thermo Fisher), and cDNAs were cloned into pCR8 by TOPO-TA cloning (Thermo Fisher) before Sanger sequencing.

### Relative copy number determination of SkV-L-A1 in *S. kudriavzevii*

*S. kudriavzevii* was transformed with plasmids expressing *XRN1* from various *Saccharomyces* species, and an empty vector control using the LiAc method. The transformation was carried out at room temperature and heat shocked at 30°C. *S. kudriavzevii* transformants were recovered on CM –tryptophan and grown at room temperature. Clones were derived from two independent transformation reactions and grown to mid-log phase at room temperature. Total RNA was extracted from these cultures by first treating the cultures with Zymolase 100T (final concentration 100 µg mL^-1^) for 2 hours at room temperature in buffer Y1 (1 M Sorbitol, 100 mM EDTA (pH 8.0), 14 mM β-mercaptoethanol). Yeast spheroplasts were treated with Trizol to extract total cellular RNA, followed by a digestion of residual DNA by Turbo DNase for 30 min at 37°C (Thermo Fisher). The RNeasy RNA cleanup protocol was used to remove DNase from the RNA samples (Qiagen), which were then stored at -80°C. RNA was converted to cDNA using Superscript III and random hexamer priming, as per manufacturers recommendations. cDNA samples were diluted 10-fold with distilled RNase-free water and used as templates for qPCR. Primers designed to recognize the RNAs corresponding to *GAG* of SkV-L-A1 (5’-TGCTTCTGATTCTTTTCCTGAATGG-3’ and 5’-GCCACTTACTCATCATCATCAAAACG-3’) and the cellular transcripts from *TAF10* (5’-ATGCAAACAATAGTCAAGCCAGAGC-3’ and
5’-TCACTGTCAGAACAACTTTGCTTGC-3′) were used to amplify cDNA using SYBR® Green PCR Master Mix (Thermo Fisher) on a CFX96 Touch^TM^ (Biorad). *TAF10* was used as a cellular reference gene to calculate the amount of viral cDNA within a given sample using the comparative *C*_T_ method [65].

## Acknowledgements

The authors would like to thank Justin Fay, Emily Feldman, David Garfinkel, Aashiq Kachroo, Maryska Kaczmarek, Lawrence Kelley, Ed Louis, Nicholas Meyerson, Susan Rozmiarek, Soumitra Sau, Alex Stabell, Nahum Sonenberg, Cody Warren, Reed Wickner, Chris Yellman and Renate van Zandwijk for critical reagents, laboratory support, and insightful discussions.

## SUPPORTING INFORMATION LEGENDS

**Figure S1.**
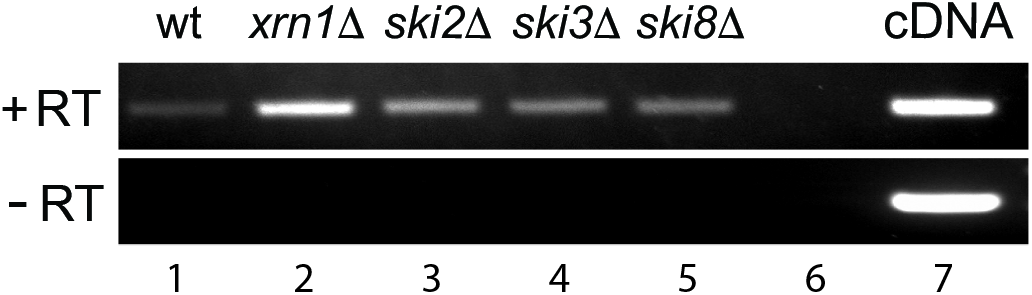
Confirmation that L-A virus dsRNA can be detected within *S. cerevisiae*. We wished to confirm that the dsRNA being detected in figure 2A of the paper was actually L-A in origin. dsRNA samples were used as templates for L-A negative-strand-specific cDNA synthesis and subsequent PCR amplification. Specificity of our primers for L-A was confirmed by a positive control where plasmid cloned L-A cDNA was used as a template (lane 7) and a negative control where RNA extracted from *S. cerevisiae* 2405 (L-A-, M-) was used as a template (lane 6).

**Figure S2.**
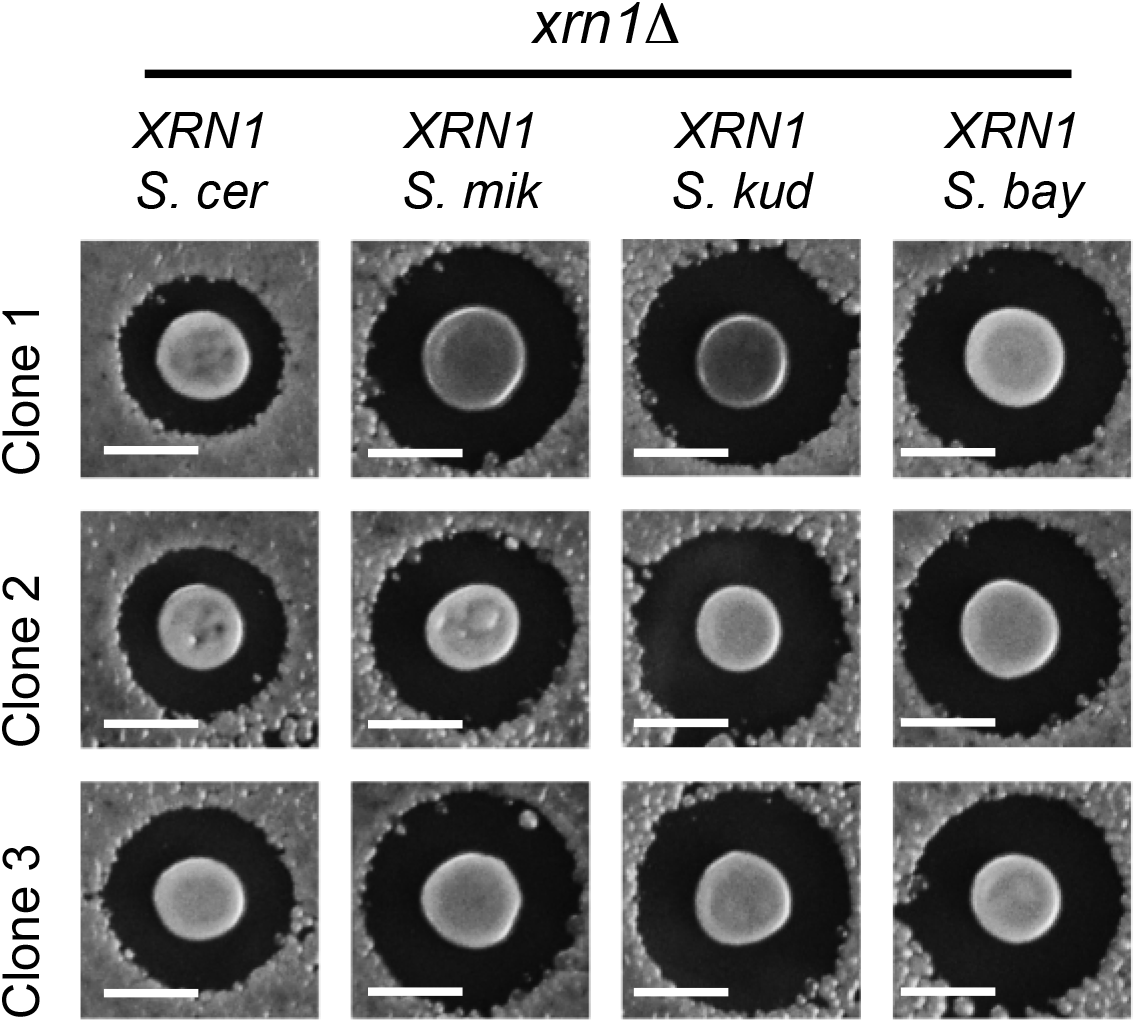
Expression of *XRN1* and its impact on killer toxin production within *S. cerevisiae*. Representative pictures of kill zones produced by individual clones of *S. cerevisiae xrn1*Δ L-A^+^ Killer^+^ expressing *XRN1* from different species. The scale bar represents 5 mm.

**Figure S3.**
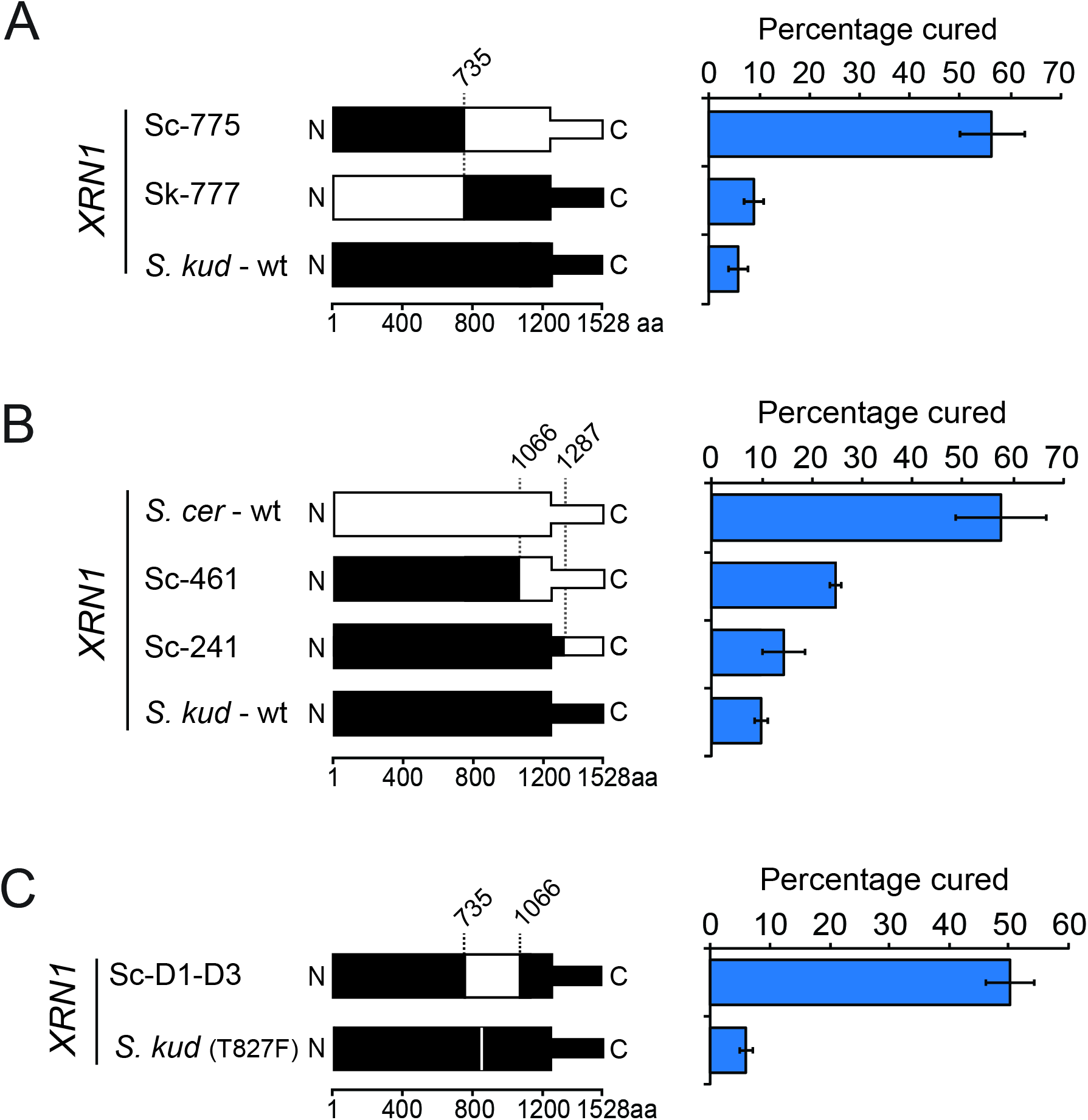
Chimeric *XRN1* genes reveal the importance of the C-terminal domain for efficient curing of the killer phenotype. (*Left*) Schematic representations of chimeric proteins derived from various C-terminal domain fusions between *XRN1* from *S. cerevisiae* (white) and *S. kudriavzevii* (black). Dotted lines represent the boundaries of the chimeric fusions with the numbering representing the amino acid position. (*Right*) Clonal isolates of a killer *S. cerevisiae* strain expressing chimeric Xrn1p proteins were assayed for loss of the killer phenotype resulting in “cured” clones (error bars represent SEM, n>3).

**Figure S4.**
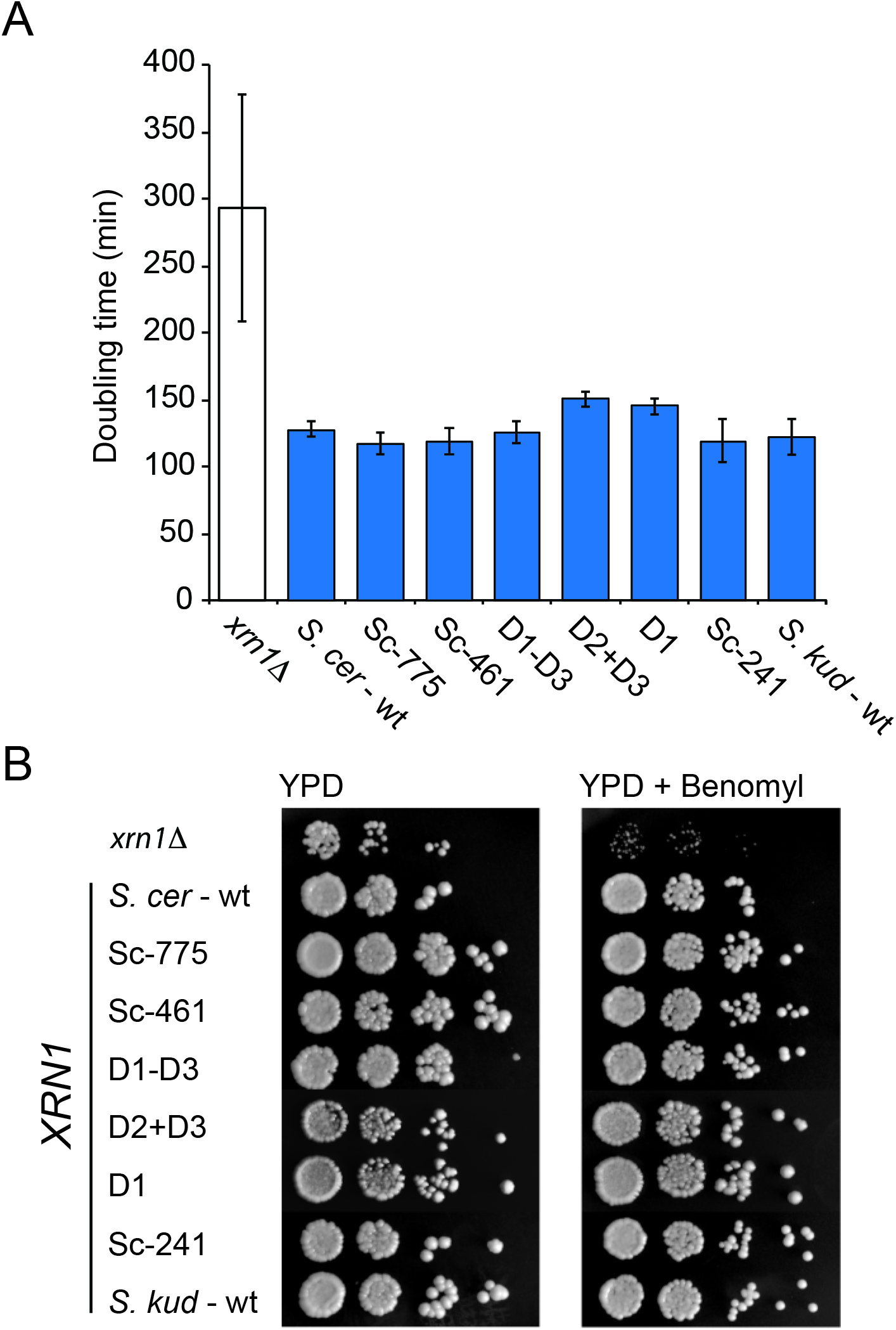
Chimeric *XRN1* genes are able to restore normal growth and benomyl resistance to *S. cerevisiae xrn1Δ*. (A) The doubling time of *S. cerevisiae xrn1* complemented with *Saccharomyces XRN1* chimeras and *XRN1* from *S. cerevisiae* and *S. kudriavzevii*. (B) The growth, morphology, and benomyl sensitivity of *S. cerevisiae xrn1Δ* cultured on YPD solid media, and the effect of complementation with *XRN1* chimeras.

**Figure S5.**
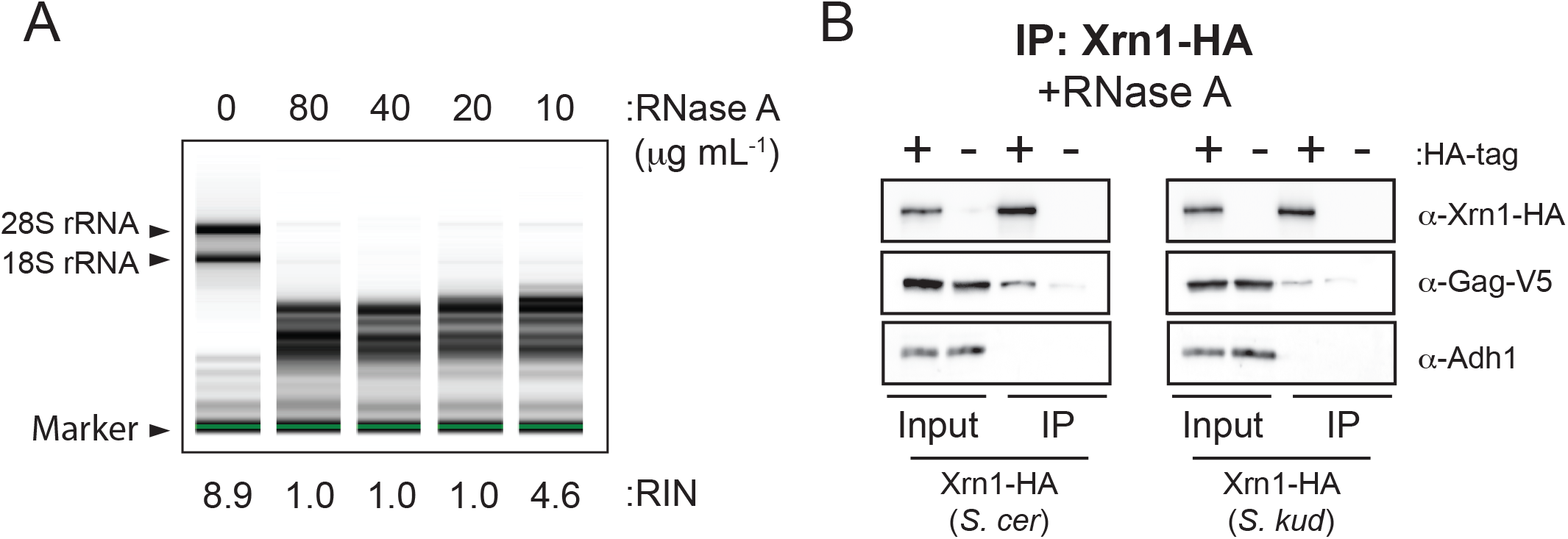
Co-immunoprecipitation of Gag by Xrn1p is not affected by the digestion of cellular RNAs. (A) The extent of RNA degradation by RNase A in yeast whole cell lysates was measured using a 2200 TapeStation Instrument with different concentrations of RNase A. RNA integrity numbers (RIN) were calculated to assess the integrity of the RNA within whole protein extract samples with and without the addition of RNase A [93]. (B) Western blot analysis of Xrn1p and L-A Gag co-immunoprecipitation. HA-tagged and untagged Xrn1p from either *S. cerevisiae* or *S. kudriavzevii* were immunoprecipitated in the presence of Gag-V5 with the addition of RNase A. Adh1 was used in all panels as a loading control to ensure equal input of total protein and the specificity of immunoprecipitation.

**Figure S6.**
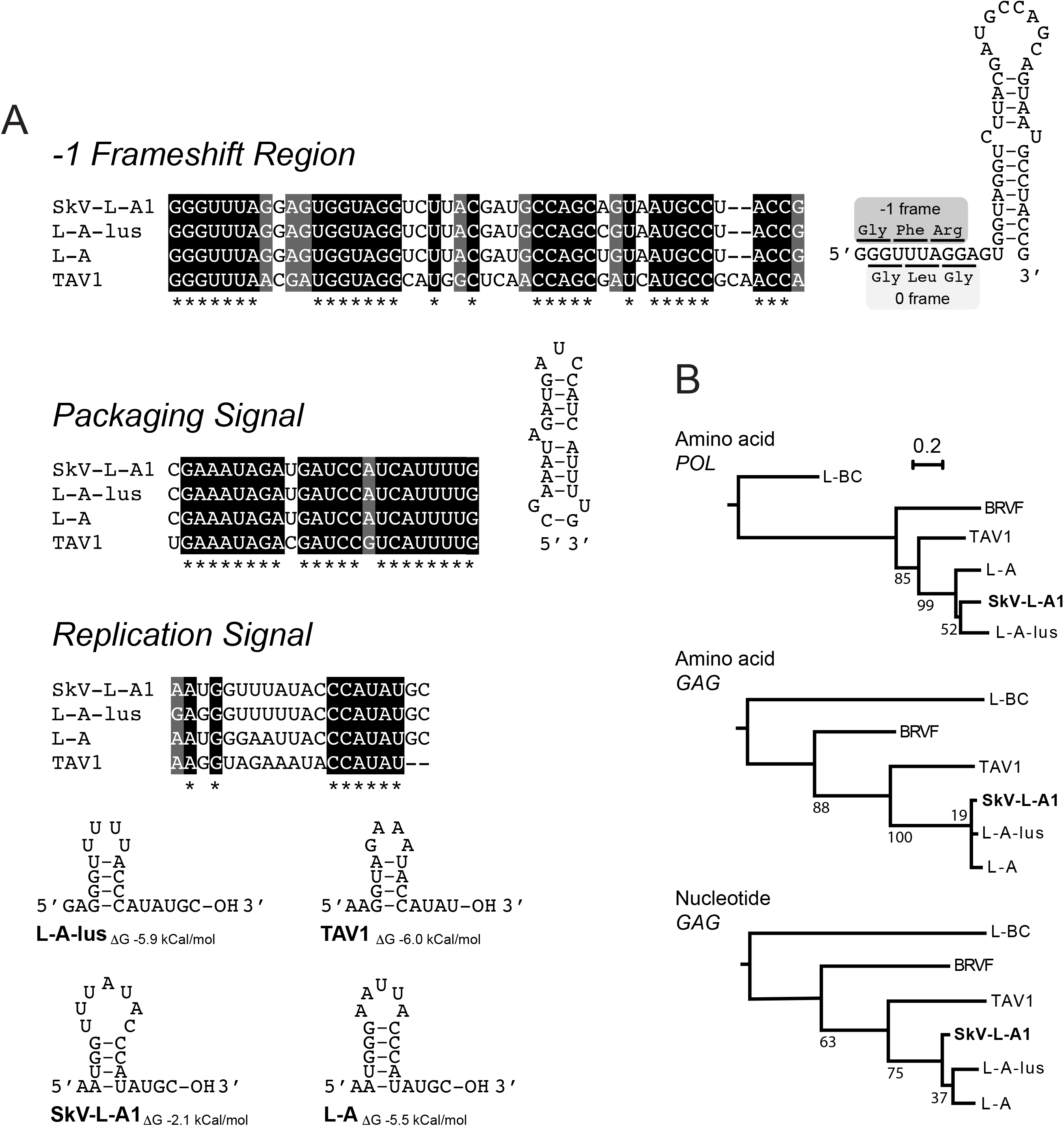
RNA sequence conservation and the phylogenetic relationship of L-A-like totivirus. (A) RNA secondary structure models of functionally important totivirus RNA sequences are based upon the sequence of SkV-L-A1 unless otherwise stated, and show the predicted base pairing between nucleotides. (B) The evolutionary history of totivirus was inferred by using the Maximum Likelihood method with bootstrap values from 100 replicates shown at each node. The amino acid and nucleotide sequence of the *POL* and *GAG* gene from six totiviruses (GenBank accession numbers: SkV-L-A1 (this study; KX601068), L-A-lus (JN819511), L-A (NC_003745), tuber aestivum virus 1 (TAV1) (HQ158596), black raspberry virus F (BRVF) (NC_009890), L-BC (NC_001641)). The tree is drawn to scale, with branch lengths measured in the number of substitutions per site.

**Figure S7.**
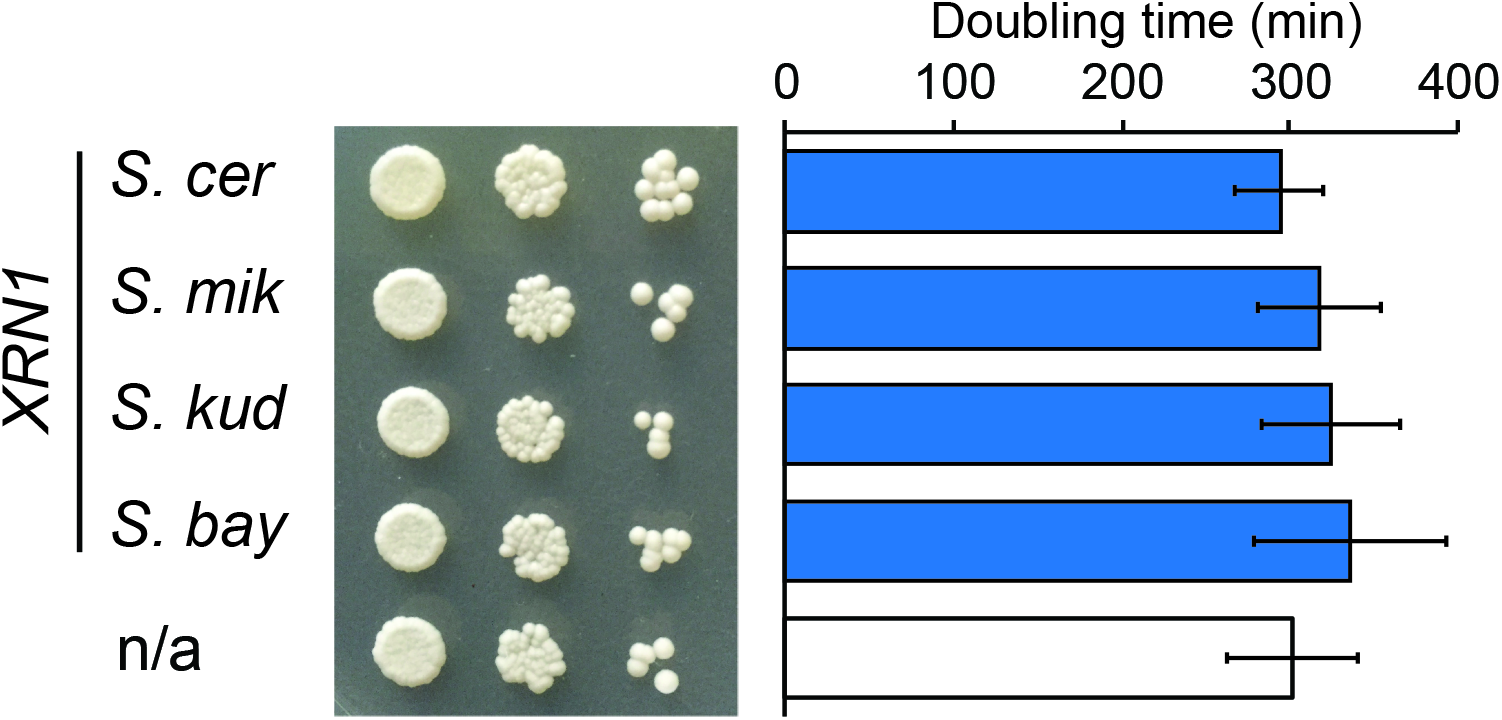
Expression of heterospecific *XRN1* within *S. kudriavzevii* and the effect on growth and colony morphology. The growth of *S. kudriavzevii* expressing each of these *XRN1* genes was measured by growing upon agar plates (left) or in liquid culture (right) and comparing to a wildtype strain that was not complemented with *XRN1*.

Table S1. **Evolutionary analysis of genes involved in RNA metabolism.** This table summarizes the results from all of the evolutionary analyses that were performed.

Table S2. **Relevant plasmids.** This table lists information on the various plasmids constructed and/or used in this study.

Table S3. **Relevant yeast strains and species.** This table lists information on the various *Saccharomyces* strains constructed and/or used in this study.

File S1.**Xrn1 structure.** This file contains PDB coordinates for the PHYRE modeled structure of *S. cerevisiae* Xrn1p.

File S2.**Plasmid sequences.** This file contains sequences of plasmids constructed as part of this study.

